# 2P-NucTag: on-demand phototagging for molecular analysis of functionally identified cortical neurons

**DOI:** 10.1101/2024.03.21.586118

**Authors:** Jingcheng Shi, Boaz Nutkovich, Dahlia Kushinsky, Bovey Y. Rao, Stephanie A. Herrlinger, Emmanouil Tsivourakis, Tiberiu S. Mihaila, Margaret E. Conde Paredes, Katayun Cohen-Kashi Malina, Cliodhna K. O’Toole, Hyun Choong Yong, Brynn M. Sanner, Angel Xie, Erdem Varol, Attila Losonczy, Ivo Spiegel

**Author notes:** Co-first authors (Jingcheng Shi, Boaz Nutkovich, Dahlia Kushinsky, Bovey Y. Rao, Stephanie A. Herrlinger). Co-senior authors (Attila Losonczy, Ivo Spiegel).

## Abstract

Neural circuits are characterized by genetically and functionally diverse cell types. A mechanistic understanding of circuit function is predicated on linking the genetic and physiological properties of individual neurons. However, it remains highly challenging to map the molecular properties onto functionally heterogeneous neuronal subtypes in mammalian cortical circuits *in vivo*. Here, we introduce a high-throughput two-photon nuclear phototagging (2P-NucTag) approach for on-demand and indelible labeling of single neurons via a photoactivatable red fluorescent protein following *in vivo* functional characterization in behaving mice. Using this novel function-forward pipeline to selectively label and transcriptionally profile previously inaccessible ‘place’ and ‘silent’ cells in the hippocampus of behaving mice, we identify unexpected differences in gene expression between these hippocampal pyramidal neurons with distinct spatial coding properties. Thus, 2P-NucTag opens a new way to uncover the molecular principles that govern the functional organization of neural circuits.

**One-Sentence Summary:** 2P-NucTag - A novel high-throughput on-demand phototagging approach to identify selective gene expression of functionally distinct neurons *in vivo* in behaving animals.

## Introduction

Information processing in neural circuits requires precise interactions between molecularly and functionally diverse populations of neurons. Since gene expression dictates neuronal connectivity and function, a fundamental goal of neuroscience has been to characterize gene expression profiles of functionally defined neurons and to measure changes in gene expression associated with distinct functional states of neurons^1^. High-throughput transcriptomic approaches such as single-cell/-nucleus RNA-sequencing (sc/snRNA-seq) and spatial transcriptomics have greatly accelerated the identification of gene programs in molecularly distinct types of neurons at single-cell resolution^2–8^. However, functional and anatomical characterization of such molecularly identified neuronal subtypes^9–13^ remains highly challenging as it requires the generation and validation of new subtype-specific molecular tools^14–16^. Recently, correlated *in vivo* Ca^2+^ imaging with *post hoc* spatial transcriptomics (ST) has been used to relate gene expression with *in vivo* function^17–20^, but this approach is limited by the relatively low density of the cells investigated and by the number of genes that can be probed in ST experiments. Furthermore, this approach is not suitable for unbiased transcriptomic analyses as selecting probes for ST requires prior knowledge of the genes of interest. Therefore, a method for unbiased identification of genes that are differentially expressed in densely packed, functionally distinct glutamatergic pyramidal neurons (PNs) would significantly accelerate our understanding of how gene expression determines circuit function and behavior.

The inability to tag single functionally identified cortical PNs *in vivo* in behaving animals presents a significant challenge as large-scale neural recordings have shown that PNs are highly heterogeneous in their physiological, anatomical, and response properties, and are spatially intermixed within neocortical and hippocampal circuits^16,21–27^. For example, PNs with distinct spatial coding properties are distributed throughout the dense cell body layer of the hippocampus^28–35^. However, the origin of this functional diversity in feature selectivity is largely unknown, and it remains unclear if gene expression differences are associated with discrete and transient functional cell states^35–40^. Thus, there is a critical need for function-forward approaches to directly identify the transcriptional profiles of cortical neurons that were functionally characterized *in vivo* in behaving animals, and to test subsequently whether the functional differences between seemingly identical neurons are indeed driven by specific differentially expressed genes.

Previous attempts using Ca^2+^ and light-dependent labeling of transiently active neurons^16,41–43^ were limited by their spatial resolution, deficiencies in targeting neurons with high baseline intracellular Ca^2+^ levels, and the inability to label neurons that decrease their activity in response to behavioral state or sensory stimuli. Similarly, labeling approaches based on the expression of immediate early genes ^44–47^ lack the temporal and spatial resolution to faithfully report the precise activity patterns and response properties of single neurons that are active but functionally distinct (e.g., neurons with different response tuning properties or feature selectivity). Finally, previous attempts to tag cortical neurons with photoactivatable fluorescent proteins^48^ with single-cell precision have been deployed with limited success^49^. Thus, although these methods, collectively, have contributed greatly to the understanding of how co-active cells contribute to behavior and cognition, there is an unmet need for an approach that allows for identifying the molecular and cellular properties of single neurons that were functionally characterized *in vivo* based on their precise activity patterns in behaving animals.

Here we introduce 2P-NucTag, a robust *in vivo* pipeline for in-depth molecular and cellular analyses of single neurons that were functionally characterized in behaving animals. The 2P-NucTag approach optimizes a previously described *ex vivo* framework^50^ and is based on the co-expression of a photoactivatable red fluorescent protein (PAmCherry) and a genetically encoded green Ca^2+^ indicator (GCaMP7f or GCaMP8s) - thereby, 2P-NucTag allows for combining large-scale *in vivo* two-photon (2P) functional imaging of cortical PNs with reliable and selective 2P phototagging of the nuclei of single neurons based on their functional properties as well as with *post-hoc* in-depth unbiased transcriptomics and cellular analyses of the photolabeled neurons. Using this novel approach, we identify unexpected differences in gene expression and cellular properties in previously inaccessible ‘place’ and ‘silent’ cells in the hippocampus of behaving mice. Thus, we achieve previously unattainable molecular characterization of functionally identified PNs *in vivo* in behaving animals using the 2P-NucTag approach.

## Results

### *In vivo* two-photon phototagging with 2P-NucTag

A major challenge in combining *in vivo* functional recording with stable tagging in the same neuron is the co-expression of an activity sensor and a photoactivatable tag with spectrally separable fluorescent imaging and photoactivation. We overcame this challenge by generating a bicistronic construct on a recombinant adeno-associated viral (rAAV) backbone that co-expresses cytosolic GCaMP7f^51^ for 2P Ca^2+^ imaging and a nucleus-targeted photoactivatable red fluorescent protein (H2B-PAmCherry) for 2P phototagging^50^ via a promoter that is selective for cortical glutamatergic neurons^52^ (2P-NucTag, Figure 1A). Upon injection of the 2P-NucTag rAAV into the CA1 region of the mouse dorsal hippocampus to label CA1 PNs, we found that GCaMP7f is properly expressed in the perinuclear space of the infected PNs (Figure 1B). Targeting the nuclei of GCaMP-expressing neurons, we achieved rapid nuclear PAmCherry photoconversion using 810-nm excitation light on a three-dimensional acousto-optical deflector microscope (3D-AOD)^53,54^ (Figure 1B; supplementary movie 1, see methods) and orthogonal GCaMP-Ca^2+^ activity imaging with 940 nm excitation light (Figure S1A-B). Photoconverted PAmCherry red fluorescence was detected at >1000 nm excitation (1040 or 1070 nm, see Methods) and was localized to the targeted nuclei (Figure 1B, top). 2P-NucTag enabled the imprinting of arbitrary tagging patterns into the CA1 pyramidal cell layer with 3D-AOD scanning, showing spatially precise photoconversion of H2B-PAmCherry with single-nuclear and even sub-nuclear resolution (Figure 1B, middle and bottom). We carried out a detailed characterization of wavelength, laser power, and duration-dependence of PAmCherry photoactivation *in vivo* to identify the optimal parameters for spatially precise photoactivation (Figure 1C). Based on our results, we opted for 810 nm excitation light with 37-42 mW laser power (measured after the objective), 1.3 ms/pixel dwell time over 70 pixel x 70 pixel regions-of-interest (ROIs, with 0.1 μm/pixel resolution). The 810-nm wavelength is spectrally separated from the GCaMP-based Ca^2+^ imaging wavelength at 940 nm, and these photoactivation wavelengths, laser power, and duration parameters yielded robust increases in PAmCherry fluorescence of targeted nuclei (192-379% increase in PAmCherry red fluorescence visualized using 1040 nm excitation: n = 36 cells, 295% ± 8% 1′F/F, mean ± s.e.m.) after single scans while minimizing total scan time and power. We confirmed that phototagged nuclei remained detectable over multiple days after photolabeling (Figure 1D). We next photoactivated a subset of CA1 PNs in a large (700 x 700 μm) field of view (FOV), similar to the FOV size used for *in vivo* 2P population imaging experiments in CA1 (Figure 1E, left). We found clear photoactivation of individual target nuclei when visualized *in vivo*. We then confirmed that *in vivo* photolabeling was preserved in *post hoc* histological slices (Figure 1E, middle), and that phototagged cells can be reliably registered across *in vivo* z-stacks and *post hoc* confocal images (Figures 1E, right, 1F, S1C-D, supplementary movie 2). In addition, we segmented the nuclei from both the *in vivo* and *ex vivo* z-stacks to observe the average axial and lateral fluorescence profiles of phototagged nuclei, demonstrating single-nucleus resolution (Figure 1F, right).

**Figure 1.**
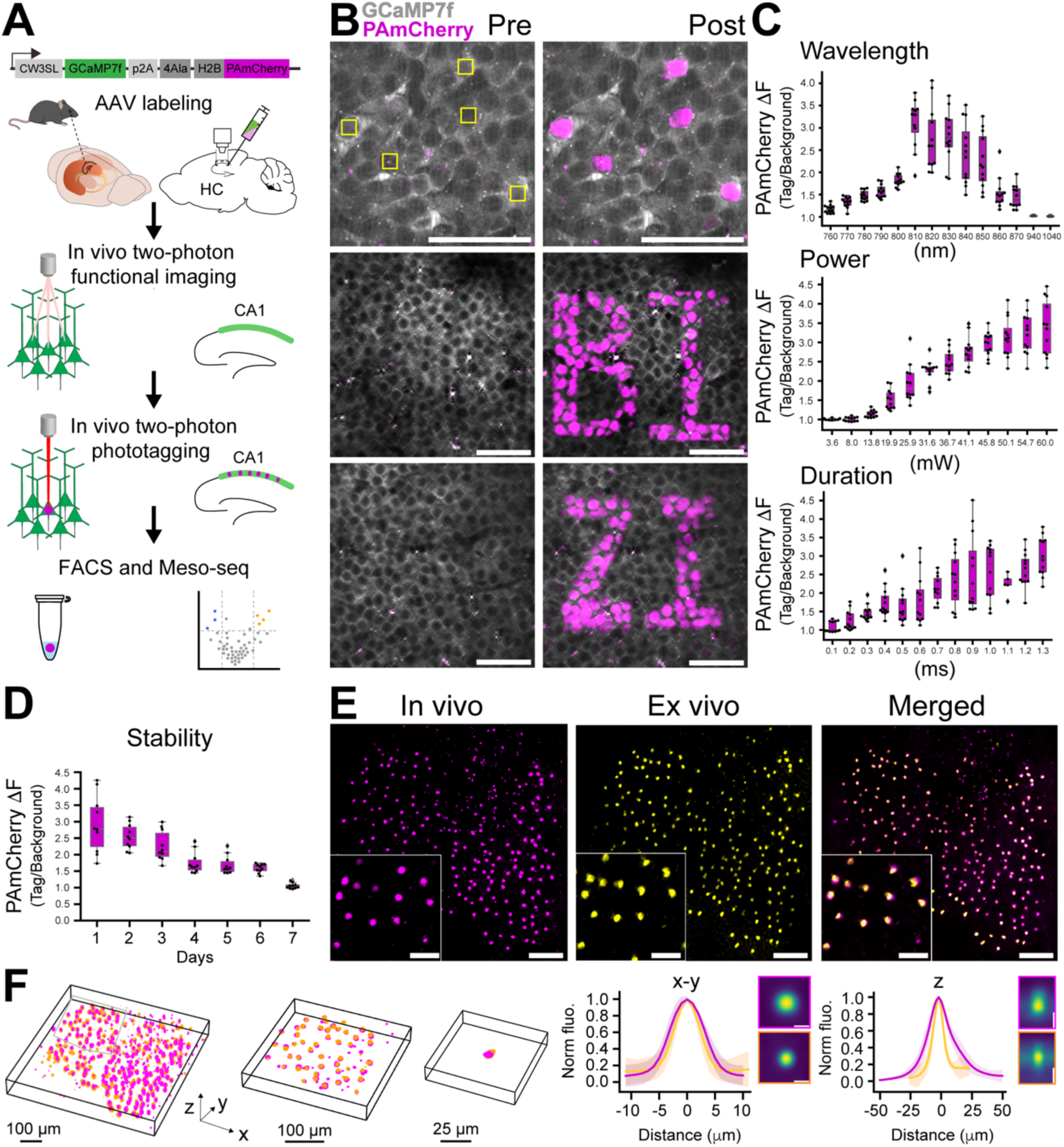
*In vivo* two-photon phototagging with 2P-NucTag. (**A**) Schematics of the 2P-NucTag pipeline. Top: bicistronic rAAV construct, injection to the hippocampus. Middle: *in vivo* two-photon (2P) GCaMP-Ca^2+^ population imaging followed by 2P PAmCherry photoactivation, fluorescence-activated cell sorting (*FACS*), and mesoscale sequencing (*Meso-seq*). (**B**) Top: representative *in vivo* time-averaged (6 frames average) 2P images of individual cells before (*Pre*) and after (*Post*) *in vivo* two-photon PAmCherry photoactivation in the CA1 pyramidal layer of the mouse dorsal hippocampus. Individual nuclei were photoactivated with 810-nm 2P laser chessboard scanning region-of-interest (ROI, yellow boxes) over target nuclei (70 x 70 pixel for each ROI, 0.1 μm/px, 1.3 ms/px total pixel dwell time, 6,370 ms total scan time per ROI. Laser power was 40 mW measured after the objective) with a 3-dimensional acousto-optical deflector microscope (3D-AOD). Gray: GCaMP7f (940 nm excitation), magenta: PAmCherry, (1040 nm excitation). Scale bar: 50 μm. Middle and bottom: imprints of letters ‘*BI*’ and *‘ZI’* following patterned *in vivo* two-photon photoactivation in the hippocampal CA1 pyramidal layer (scale bar, 50 μm). (**C**) Characterization of *in vivo* 2P photoactivation parameters for PAmCherry: duration, wavelength, laser power (measured after the objective) (n = 11-12 cells per condition). Relative change in PAmCherry red fluorescence (ΔF) is based on normalizing the tagged nuclei fluorescence to the fluorescence of neighboring untagged nuclei measured with 1040 nm excitation. **(D**) *In vivo* stability of the PAmCherry fluorescence signal over days after a single photoactivation scan (n = 8 cells). (**E**) Representative time-averaged images from z-stacks of photoactivated nuclei *in vivo* (magenta: PAmCherry, scale bar: 100 μm). Middle: *ex vivo post hoc* confocal z-stack image of the same field of view (FOV, magenta: PAmCherry). Right: registered *in vivo* and *ex vivo* images following non-rigid image transformation (magenta: *in vivo*, yellow: *ex vivo*, see methods). (**F**) Left: 3D overlay of tagged nuclei registered between *in vivo* (magenta) and *ex vivo* (yellow) z-stacks with increasing lateral resolution (as in *E*). Gray box represents the segmented area for subsequent images. Right: normalized lateral (*x-y*, left) and axial (*z*, right) fluorescence profiles (mean ± s.e.m.) of tagged cells *in vivo* (magenta, n = 1 mouse, 200 cells). Yellow: mean ± s.e.m. of *ex vivo* confocal images (as in *E* and *F,* same mouse and nuclei). Inset: (x-y) top: average *in vivo* maximum z–projection, bottom: average *ex vivo* maximum z projection; (z) top: average *in vivo* lateral projection, bottom: average *ex vivo* lateral projection. Scale bar: 10 μm. Boxplots show the 25^th^, 50^th^ (median), and 75^th^ quartile ranges, with the whiskers extending to 1.5 interquartile ranges below or above the 25^th^ or 75^th^ quartiles, respectively. Outliers are defined as values extending beyond the whisker ranges.

To establish the compatibility of the 2P-NucTag construct with downstream cell sorting and transcriptomic applications, we further prepared CA1 PN samples and subjected them to FACS and Meso-seq (Figure S2). We chose to use our previously developed Meso-seq protocol due to its advantage of providing a robust and cost-effective method for sequencing low numbers of input nuclei. We collected 3 samples from AAV-transduced nuclei (‘Injected, transduced’), 2 samples of nuclei that were not transduced but were isolated from the mice that received AAV injections (‘Injected, non-transduced’), and 2 negative control samples from mice that were not injected with AAVs (‘non-injected’). For each sample, 50-100 nuclei were collected and analyzed in bulk via Meso-seq. Between these groups, we found that they have comparable sequencing statistics in terms of total reads and percent of uniquely mapped reads (Figure S2A); notably, these sequencing statistics were similar to those when we applied Meso-seq to AAV-infected visual cortex samples in previous studies^55,56^. Gene expression levels between groups were highly correlated (Figure S2B-C) and reads of representative genes were similar between groups except for the inhibitory neuron markers (Figure S2D). Thus, our results demonstrate that the previously established Meso-seq protocol works well with low numbers of hippocampal PN nuclei as input, and that the 2P-NucTag construct does not cause changes in transcriptional properties in AAV-infected PNs. We also obtained *ex vivo* whole-cell patch-clamp intracellular recordings from phototagged and control CA1 PNs (Figure S3A) in acute hippocampal slices: comparing H2B-TAG (infected and photoactivated), H2B (infected), and control (non-infected) cells revealed no differences in intrinsic properties (Figure S3B-G). Together, these results confirm the utility of 2P-NucTag to analyze *ex vivo* the cellular properties of single CA1 PNs that were functionally characterized and phototagged *in vivo* and demonstrate that the 2P-NucTag construct does not affect the intrinsic physiological properties of these PNs. Thus, 2P-NucTag enables high-throughput, indelible phototagging of neuronal nuclei *in vivo* that can also be identified via our registration pipeline for *post hoc* analyses *ex vivo*. Phototagged nuclei further enable downstream RNA-sequencing and *ex vivo* electrophysiology analyses.

### Phototagging of functionally identified PNs in the hippocampus *in vivo*

To demonstrate the utility of 2P-NucTag for on-demand labeling of functionally defined PNs, we deployed 2P-NucTag to label hippocampal PNs located in the dorsal CA1, a region with well-established spatial coding heterogeneity^28–35^. Only a subset of hippocampal PNs (‘place cells’) exhibit reliable spatial tuning for a location (‘place field’) during exploration^57^. Place cell identity of PNs^58^ is highly dynamic, with the majority of cells changing their spatial tuning properties over the timescale of days^35–40^. While place cell properties of PNs were thought to be randomly allocated onto seemingly homogeneous PNs^58^, recent studies demonstrated that the genetic expression profiles of CA1 PNs are also highly heterogeneous^30,32,59^. These transcriptional differences are present along all anatomical axes of the hippocampus^29^, manifest at the protein expression level, and are comprised of genes with known functions (e.g., transcription factors, cell adhesion molecules, voltage-gated channels, neurotransmitter receptors, and auxiliary subunits)^29,60^. Gene expression differences thus could produce variability in the biophysical properties and connectivity of PNs and may underlie the differences in stable spatial coding properties of CA1 PN subpopulations. Thus, the spatial intermingling of place cells with active-non-place and silent PNs^61–64^ without apparent topographical organization in the densely packed CA1 pyramidal layer^35,64^ allows us to test the utility of 2P-NucTag in selectively labeling PNs occupying these distinct functional states.

To selectively label CA1 PNs with distinct spatial coding properties, we trained mice in a spatial navigation task for water rewards in a linear virtual reality environment^65–67^ and performed *in vivo* 2P GCaMP-Ca^2+^ imaging of PNs (Figure 2A-B). We reliably detected GCaMP-Ca^2+^ signals from individual PNs (Figures 2C, S1B). We analyzed the basic characteristics of GCaMP7f in the bicistronic 2P-NucTag construct and confirmed that the basic properties of the indicator are similar to those for a previously published single-construct version^51^ of GCaMP7f (Figure S1B). We classified all PNs in the imaging FOV as place cells, active non-place cells, or silent cells (Figure 2D, see methods). Place cells were identified as described in our previous studies^38,67–69^ where place fields were detected by identifying spatial tuning curves surpassing the 95^th^ percentile of shuffled tuning curve values, and silent cells were classified as cells with no deconvolved events in the recording session. Following the functional identification of PNs, we generated spatial masks of place cell locations in the FOV and used these masks to guide 2P phototagging of all identified place cells in the imaging FOV with 3D-AOD (n = 5 mice, Figure 2E-F). In a separate set of mice (n = 4 mice), we tagged a subset of CA1 PNs that exhibited no detectable activity during the imaging session (‘silent cells’ see methods, Figure S4, Table 1), whereby we photoactivated a similar number of silent cells as place cells in each mouse despite their higher abundance in our recordings. We could reliably register photoactivated nuclei to the spatial masks of functional profiles that we generated for cells of interest (Figures 2F, S4D-E, S5, see methods). We quantified the number of place cell nuclei that we successfully photoactivated and compared it to the number of spatial masks we generated for each animal. For silent cells, we performed a similar calculation with number-matched silent cell nuclei. Across all animals, we phototagged 93.3% ± 4.2% (n = 5 mice) of all place cell masks and 53.6 ± 6.6% (n = 4 mice) of all silent cells (Figure 2F).

**Figure 2.**
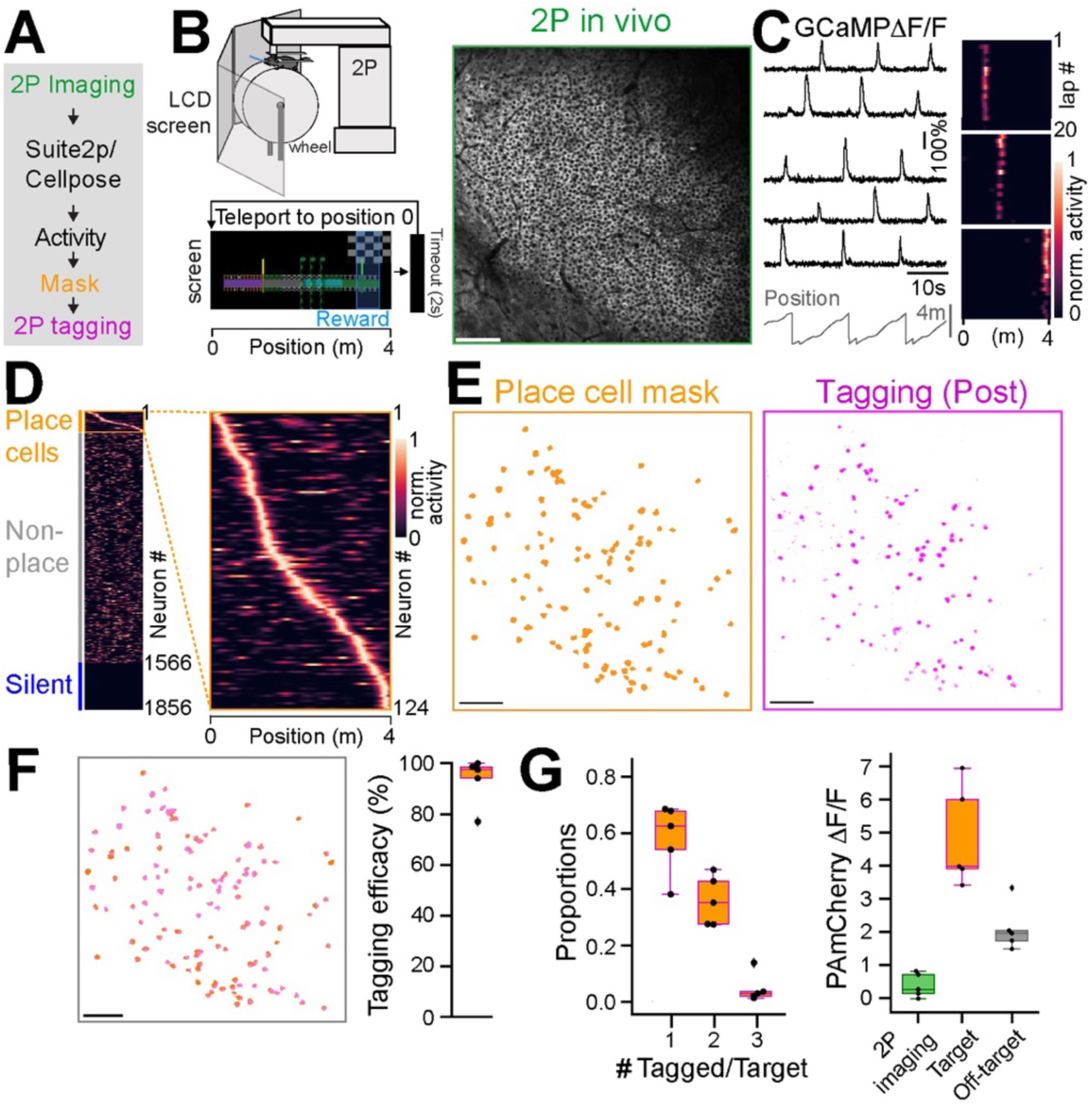
Selective phototagging of place cells in the hippocampus with 2P-NucTag. (**A**) Pipeline for two-photon (*2P*) phototagging of functionally identified hippocampal neurons during spatial navigation. (**B**) Left: schematics of 2P imaging setup in virtual reality (VR). Head-fixed mice are trained to run for a water reward in a 4-m long linear VR corridor projected onto LCD screens surrounding the animal. At the end of the corridor, mice are teleported back to the start position after a 2-second delay. Right: example 2P field of view (FOV) of GcaMP in the CA1 pyramidal layer. Scale bar: 100 μm.(**C**) Left: Traces of relative GcaMP-Ca^2+^ fluorescence changes (ΔF/F) from five example CA1 place cells during VR spatial navigation. Right: heatmaps of normalized ΔF/F activity from three example place cells over 20 laps during VR navigation. (**D**) Left: heatmap of all CA1PNs detected with Suite2p/Cellpose in the FOV shown in *B*. Identified place cells are marked with an orange box. Right: Zoomed-in heatmap of place cell tuning curves. (**E**) Left: spatial mask (orange) of identified place cells from *D* in the FOV. Right: PAmCherry fluorescence (magenta) of tagged nuclei after 2P phototagging. Scale bar: 100 μm. (**F**) Left: overlay of spatial masks of identified CA1PNs and tagged nuclei for the FOVs in *E*. Scale bar: 100 μm. Right: tagging efficacy, defined as the fraction of successfully tagged place cell nuclei (93.3% ± 4.2%, mean ± s.e.m., n = 5 mice). (**G**) Left: Proportion of single, double, and triple-tagged nuclei following phototagging of a single place cell. Right: relative change in PAmCherry red fluorescence (1070 nm excitation) for non-tagged cells in the FOV after 2P imaging (green), after 2P phototagging of targeted place cell nuclei (orange) and off-target nuclei (gray, n = 5 mice). Boxplots show the 25^th^, 50^th^ (median), and 75^th^ quartile ranges, with the whiskers extending to 1.5 interquartile ranges below or above the 25^th^ or 75^th^ quartiles, respectively. Outliers are defined as values extending beyond the whisker ranges.

**Table 1.**
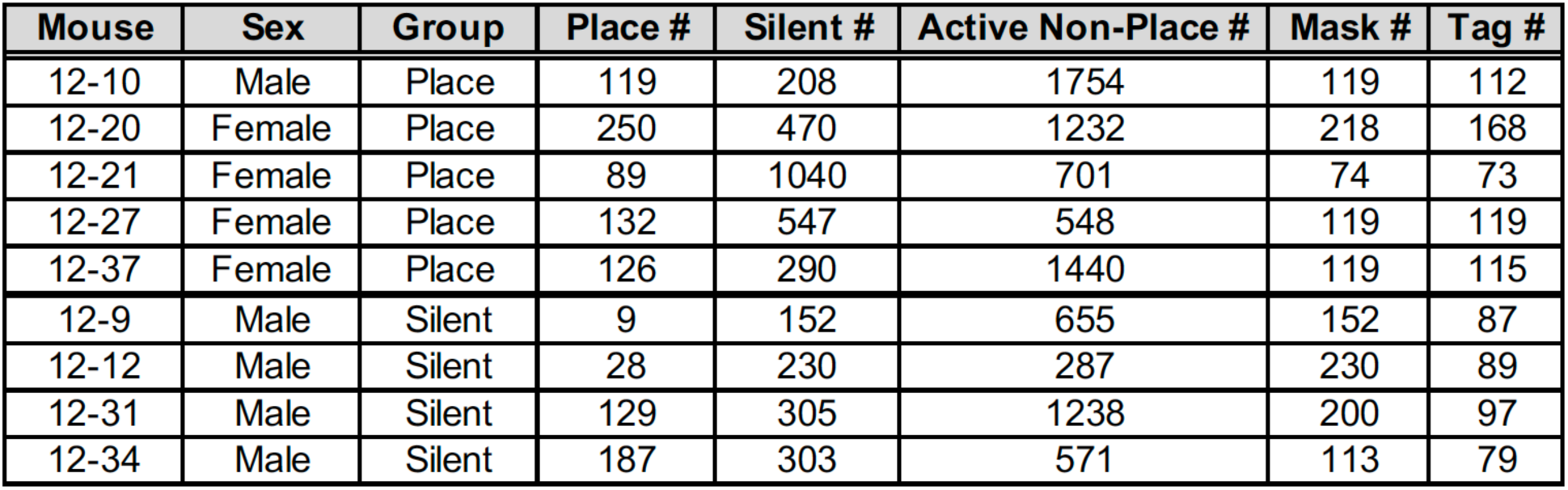

To assess the accuracy of 2P photoactivation in the densely packed CA1 pyramidal layer, we quantified the number of tagged nuclei for each targeted cell and found that photoactivation is well restricted to the target cells’ nuclei with limited off-target labeled nuclei, which had a smaller increase in mCherry fluorescence (Figures 2G, S4F). After tagging, mCherry fluorescence increased in targeted place cells (487.7% ± 67% 1′F/F, n = 5 mice) more than double compared to the background and off-target fluorescence (210.4% ± 32.1% 1′F/F, n = 5 mice). Furthermore, 2P GCaMP-Ca^2+^ imaging at 940 nm over the course of the imaging session (18-40 min, see Methods) resulted in a minimal increase of mCherry red fluorescence (37.4% ± 16.2% 1′F/F, n = 5 mice, Figure 2G, right, Figure S4F, 1070 nm excitation) and did not affect our ability to identify phototagged neurons. Finally, mouse behavior and the quality of GCaMP recordings were consistent between the two groups of mice (silent and place, Figure S4G-I).

To further confirm that photoactivation of the PAmCherry does not change the functional response properties of neurons, we expressed the 2P-NucTag construct in the primary visual cortex (V1) of adult mice and recorded visually evoked responses of neurons in this region before and after photoactivation (Figure S6A). We found that the orientation tuning (including both preference and selectivity) remained largely the same in visually responsive neurons following photoactivation (Figure S6B-C), indicating that activation of the PAmCherry does not alter neuronal response properties. These experiments were conducted in the visual cortex due to the known stability of its tuning properties^70^.

Thus, taken together, our experiments demonstrate that 2P-NucTag is suitable for on-demand *in vivo* labeling of functionally defined neurons with high efficacy and accuracy.

### Transcriptional profiling of functionally identified CA1 PNs

The ability to tag single cells *in vivo* enables a powerful new form of hypothesis generation and testing by which functionally defined cells with known behavioral relevance can be isolated and characterized. To demonstrate this, we sought to interrogate transcriptomic signatures of the functional ‘place’ and ‘silent’ cell states of CA1 PN.

Following functional imaging and *in vivo* phototagging, brain tissue containing dorsal CA1 was collected at least 25 hours after functional imaging to eliminate the impact of immediate early genes, and nuclei were dissociated and stained with a CoraLite 488-conjugated NeuN antibody and DAPI to identify neuronal nuclei (see Methods). *In vivo* photoactivated nuclei were identified by bright mCherry fluorescence (Figure 3A,B) and were isolated from the non-photolabeled neuronal nuclei by fluorescent-activated cell sorting (FACS), enabling an estimated 18-67% recovery of all photoactivated nuclei identified during in vivo imaging (Figure 3C). To validate that the FACS events with bright mCherry fluorescence were indeed the desired nuclei, we also performed sorting on samples derived from mice that were injected but not imaged; injected and imaged; or injected, imaged, and phototagged (Figure S7): we set the gates for mCherry fluorescence based on these samples and demonstrated that low mCherry fluorescence was induced by the imaging laser (Figure S7B) since the phototagged nuclei exhibited clearly identifiable higher mCherry fluorescence as compared to either of the other samples (Figure S7C). We then collected these bright nuclei collected for downstream transcriptomic analyses. To determine whether place cells and silent cells differed in their gene expression programs, RNA-seq was performed on both populations of sorted nuclei by Meso-seq, an approach that enables reliable identification of differentially expressed genes in ultra-low amounts for FACS-isolated neuronal nuclei (i.e., tens of sorted nuclei per animal, Figure 3D)^55^. After isolating the phototagged nuclei, libraries were generated with the Meso-seq protocol and were sequenced at a depth of 40-60 million reads per library (Figure S8A). Reads were aligned to the mm39 mouse genome assembly with STAR, counted with HTSeq, and gene expression patterns were compared between nuclei isolated from mice in which place cells were tagged (n = 5) and mice in which silent cells were tagged (n = 4) via PyDESeq2.

**Figure 3.**
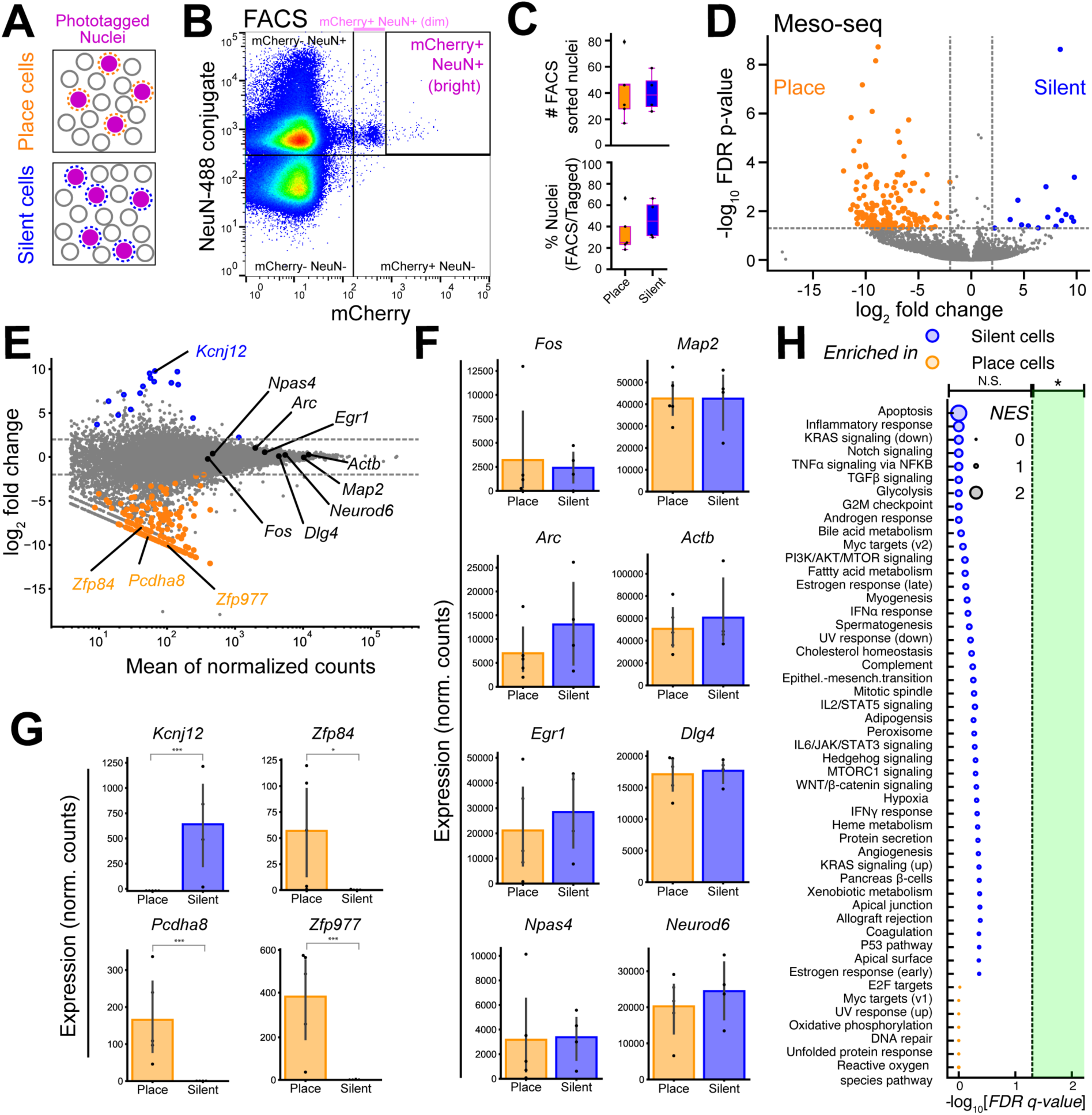
*Post hoc* transcriptional profiling of phototagged place and silent cells. (**A**) Schematics of *in vivo* photoactivated nuclei. ‘Place’ cell sample and ‘Silent’ cell samples from different mice were collected for FACS and Meso-seq. (**B**) Representative FACS graph. Gating for mCherry was set after the first 5000 events of DAPI+ nuclei to the border of the ‘dim’ mCherry+ population to separate out the sparse and high-intensity mCherry+ population. Bright mCherry+ NeuN+ populations were collected as the photoactivated nuclei. (**C**) Top: number of FACS sorted nuclei from ‘place’ and ‘silent’ samples (n = 9, 17-79 sorted nuclei, 40.3 ± 6.43, mean ± s.e.m.). Bottom: Proportion of FACS sorted nuclei compared to the number of *in vivo* photoactivated nuclei (n = 9, 18.45% to 66.39% FACS recovery, 39.94 ± 6.30%, mean ± s.e.m., ‘place’ mean = 34.63%, ‘place’ median = 25.0%; ‘silent’ mean = 46.59%, silent median = 45.09%). (**D**) Volcano plot of Meso-seq differential expressed gene (DEG) analysis for ‘place’ and ‘silent’ cells (significantly different genes are shown in orange and blue. Orange: enriched in place cells; blue: enriched in silent cells). (**E**) Meso-seq MA plot depicting DeSeq2 normalized gene counts versus log2 fold change of silent/place samples. Genes that are significantly different are labeled in orange and blue (same as above). Genes shown in panels *F* & *G* are highlighted and labeled in *E*. (**F**) Bar graph showing the normalized counts for genes that are not differentially expressed (FDR adjusted q-value. *<0.05, \*\***<**0.001, ***<0.001, PyDeSeq2. Otherwise, comparisons are not significant). (**G**) Bar graph showing the normalized counts for differentially expressed genes (FDR adjusted p-value. *<0.05, \*\***<**0.001,***<0.001, PyDeSeq2. Otherwise, comparisons are not significant). (**H**) Gene ontology analysis performed on all differentially expressed genes. Vertical line: FDR-adjusted p value of 0.05. NES = normalized enrichment score. Boxplots show the 25^th^, 50^th^ (median), and 75^th^ quartile ranges, with the whiskers extending to 1.5 interquartile ranges below or above the 25^th^ or 75^th^ quartiles, respectively. Outliers are defined as values extending beyond the whisker ranges.

In both populations, canonical CA1 PN marker genes (*Map2, Actb, Dlg4, Neurod6*) were highly and non-differentially expressed, supporting the cellular precision of our tagging approach (Figure 3E-F). However, 219 genes were identified as differentially expressed in a significant manner between silent and place cells (Figure 3E, see Supplementary Table). In some cases, specific genes were reliably identified in one group and absent from the other. For example, no counts were measured for an inward rectifying potassium channel (*Kcnj12*) in place cell samples, but the expression was present in all silent cell samples. Conversely, mRNA from two transcription factors of the zinc finger protein family (*Zfp84, Zfp977*) and a protocadherin gene (*Pcdha8*) were identified in every place cell sample and in none of the silent cell samples (Figure 3G). To ensure that the observed differences did not arise from other sources of variation between our place and silent cell samples, we performed several additional analyses, investigating potential contributions from anatomical positioning, transcriptional responses to neural activity, cellular health, and sex. Genes for which expression has been shown to vary along the dorsoventral and proximodistal axes of CA1 were not differentially expressed in our place and silent-cell nuclei (Figure S8B)^29,71^, indicating that any observed differences are unlikely to have originated from differences in anatomical positioning during cell tagging. Immediate early gene levels (*Fos, Arc, Egr1, Npas4*) did not differ (Figure 3F)^72–74^, suggesting that the ≥25 hours between the last behavior session and tissue collection (see Methods) was sufficient time to eliminate the impact of immediate transcriptional responses to neural activity. Apoptotic gene counts also did not differ between the two groups (Figure S8), indicating that there was no significant difference in cell health between the silent and place cells^75^. Finally, to test for gross molecular differences between the cells in an unbiased fashion, we performed gene set enrichment analysis on fifty hallmark gene sets from the mouse molecular signatures database and found no significant differences between silent and place cell samples (Figure 3H). Since the sex of mice used for both groups was not well balanced in this study (Table 1), it is possible that gene expression differences originated from the mouse’s sex rather than the functional identity of the tagged cells. To explore this possibility, we compared gene expression in randomly tagged (function-blind) CA1 PN nuclei from both male (n = 2) and female (n = 2) mice. Although we found that known sex-specific genes^76^ were differentially expressed, none of the genes identified as enriched in either silent or place cells had significant sex associations (Figure S9C,D). To assess if our differentially expressed genes-of-interest signified upregulation or downregulation in place cells from the mean or, conversely, downregulation or upregulation in silent cells, we compared gene expression in place cells and silent cells to randomly tagged cells separately. We found that *Kcnj12* expression was enriched in silent cells compared to random cells. *Zfp84* and *Zfp977* were significantly downregulated in silent cells compared to random cells, while *Pcdha8* expression was significantly enriched in place cells compared to random cells (Figure S9A,B).

Encouraged by our success in establishing 2P-NucTag for in-depth transcriptomic analyses in ‘place’ and ‘silent’ cells, we next sought to further develop this approach for additional key neurobiological applications. Specifically, we sought to test whether (1) the transcriptomic data obtained with the 2P-NucTag can be validated and used for dissecting the molecular mechanisms that determine functional properties of the phototagged neurons, and (2) 2P-NucTag can be used for more nuanced cell populations and resolve genetic changes of the same functional cell group at different timepoints.

For in-depth validation and functional analyses, we focused on *Kcnj12,* which our 2P-NucTag based transcriptomic analyses identified as expressed in silent cells but not in place cells. Since this gene encodes for an inward rectifying potassium channel Kir2.2^77–79^, we hypothesized that its expression could reduce the intrinsic excitability of the neurons and make them more likely to be silent. Thus, we first validated the differential expression of *Kcnj12* by performing RNAScope Fluorescent in situ hybridization (FISH) in brain sections with tagged silent CA1 PNs (Figure S10). Using a probe set for detecting *Kcnj12* (Figure S10A), we quantified the number of *Kcnj12* puncta in tagged silent cells and a number-matched subset of randomly selected non-tagged cells (n = 247 cells each, n = 3 mice). These analyses confirmed that tagged silent cells indeed express more *Kcnj12* transcripts than non-tagged cells (Figure S10B). Similar RNAscope FISH experiments for *Pcdha8* (Figure S10C) further confirmed the specificity of our 2P-NucTag-based transcriptomic analyses since *Pcdha8* expression was enriched in tagged place cells as compared to randomly selected non-tagged cells (Figure S10D, n = 321 cells in each subgroup, n = 3 mice). Next, we tested whether silent CA1 PNs differ in their intrinsic electrophysiological properties. For this, we performed targeted whole-cell patch-clamp recordings from tagged silent and nontagged cells *ex vivo* in acute hippocampal slices (Figure S11). Consistent with our hypothesis, we found that silent cells had a hyperpolarized resting membrane potential as compared to random non-tagged cells (Figure S11A). These experiments demonstrate the utility of the 2P-NucTag approach not only for molecular and analyses in functionally defined neurons but for connect *ex vivo* electrophysiological and cellular analyses of such *in vivo* labelled neurons.

Finally, to test whether the expression of Kir2.2 affects the likelihood of CA1 PNs to become place cells, we designed an AAV vector for Cre-dependent expression of Kir2.2 along with fluorescent protein mRuby3 (pAAV-hSyn-FLEX-Kir2.2_WT_-T2A-mRuby3, Figure S12E); as a negative control, we also designed a vector that drives expression of an inactive (non-conducting) mutant of Kir2.2 (pAAV-hSyn-FLEX-Kir2.2_Mut_-T2A-mRuby3, Figure S12E). We first co-injected these vectors along with an AAV to drive the expression Cre-recombinase (pAAV-hSyn-Cre) into the hippocampus of wild-type mice and assessed how ectopic expression of Kir2.2 affects the excitability of infected neurons. Using targeted whole-cell patch clamping, we found reduced excitability in cells expressing Kir2.2_WT_ (Figure S12A-D) as these cells had lower resting membrane potentials, lower input resistance and lower firing rates across current steps when compared to cells with the control virus (Figure S12B-D). We then sought to test the effect of ectopic Kir2.2 expression on functional properties of CA1 PNs *in vivo* in behaving animals. We used two sets of mice in which we co-injected either of the Kir2.2 vectors along with AAVs for neuronal expression of GCaMP8s (pAAV-hSyn-GCaMP8s) and Cre-recombinase (pAAV-hSyn-Cre) and trained mice in the same behavior paradigm as described in Figure 2. We found that the cells expressing Kir2.2_WT_ are more silent than mRuby3-negative cells in the same field of view, while these effects were not observed in cells with Kir2.2_Mut_ (Figure S12F). Comparisons of spike rate and place cell percentage further indicated a silencing of activity and a lower proportion of place cells in Kir2.2_WT_-expressing CA1 PNs (Figure S12G-H). Therefore, expression of Kir2.2 in CA1 PNs silences neurons, thereby affecting their *in vivo* function.

To further assess whether the 2P-NucTag approach can be used for molecular analyses in cells with even more nuanced differences in *in vivo* activity and function, we transcriptomically profiled place cells at different time points since a subset of these cells stabilizes their spatial representations over time and thus might differ in gene expression from when place cells first emerge. For this, we co-injected a version of 2P-NucTag AAV (pAAV-Ef1a-H2B-PAmCherry) with an AAV for pan-neuronal expression of GCaMP8s (pAAV-hSyn-GCaMP8s) into the hippocampal CA1 of two sets of mice. The mice were subsequently trained in a blank featureless environment and then recorded in a feature-rich environment (Figure S13A). In one group, we photo-tagged and collected place cells from day 0 of recording (‘Day 0’ cells), while in the other group we continuously recorded the same FOV for 5 days, and photo-tagged and collected place cells with at least 4 place fields across the 5 days of recording (‘Recurring’ cells, Figure S13B-C). Transcriptomic analyses were done with Meso-seq as described before. Using this approach, we identified 6 genes that were differently expressed (Figure S13D-F), including *Prkcd* and *Adamts1,* two genes enriched in ‘Recurring cells’ which were previously found to play a role in memory and synaptic connectivity^80,81^ and which therefore might control the synaptic wiring of neurons to maintain long-term representation. Thus, we conclude that 2P-NucTag is useful for molecular analyses in more nuanced cell types and cells with the same functional identity at different timepoints.

Taken together, our results demonstrate that the 2P-NucTag approach allows for identifying the molecular and cellular properties of neurons in the same brain region that differ only by their activity patterns *in vivo* in behaving animals. Using this approach, we identify previously unappreciated transcriptomic differences in “place” and “silent” cells in the hippocampus of behaving mice and correlate these molecular profiles with the distinct electrophysiological properties of these cells. We further revealed that these differentially expressed genes affect the function of the respective groups of neurons by validating and demonstrating the role of *Kcnj12* in the excitability of CA1 PNs and the functional consequences when ectopically expressed. The 2P-NucTag approach has the advantage of revealing previously unknown molecular profiles of PNs and works with other methods to dissect the contribution of specific molecular processes.

## Discussion

The comprehensive molecular-functional characterization of cortical circuits remains elusive due to the limited toolkit for correlated *in vivo* functional recording techniques and *post hoc* molecular analyses. Here, we introduced 2P-NucTag, a robust function-forward approach combining *in vivo* imaging, on-demand single-nucleus tagging, and *post hoc* transcriptomics. This pipeline seamlessly integrates 2P Ca^2+^ imaging in behaving mice, 2P phototagging of functionally identified neuronal nuclei, FACS-isolation of the tagged nuclei, and subsequent analyses of isolated nuclei with a mesoscale protocol for in-depth transcriptomics in ultra-sparse neuronal populations.

We demonstrated the utility of 2P-NucTag by selectively labeling and analyzing transcriptional profiles of two functionally orthogonal subpopulations of hippocampal PNs, ‘place’ cells and ‘silent’ cells, which remained inaccessible with existing techniques. Unexpectedly, our experiments revealed a number of genes that are putatively differentially expressed between these distinctly transient physiological cell states^35–40^. The relatively low expression levels for some of these genes indicate that further replicates may be necessary to confirm their differential expression. Nevertheless, several of the identified genes generate new hypotheses about the molecular mechanisms that underlie these two functional cell identities, which had been molecularly indistinguishable otherwise. Specifically, the marked enrichment of an inward rectifying potassium channel Kir2.2 (encoded by *Kcnj12*) in silent cells suggested that expression of this gene might reduce the excitability of hippocampal pyramidal neurons to thereby lower the activity of these cells *in vivo* and reduce their propensity to become place cells^82^. Indeed, our experiments based on ectopic expression of *Kcnj12* are consistent with this idea, and thereby implicate intrinsic excitability as a putative cellular and molecular mechanism that can influence the activity and spatial coding properties of PNs in the hippocampus. Thus, our findings on *Kcnj12* align with previous studies showing that intrinsic excitability is a major factor governing cell recruitment to a memory trace^83–86^. Future experiments will be needed to test the function of specific genes that we have identified with our tagging technique, which could reveal a non-random allocation of functional activity onto CA1 PNs during learning and experience. In particular, future application of 2P-NucTag to selectively tag PNs with highly distinct and shared coding properties, including place field propensity^28,36,87,88^, recruitment to ‘replay’ and ‘preplay’ events^89–92^ and participation in functionally interconnected subnetworks^69^ may uncover genetic and developmental^93–98^ backbones of functional heterogeneity among PNs.

Our results showed low levels of unintended PAmCherry photoactivation during functional imaging as well as limited off-target labeling. Although the activation level of these background and off-target neurons is lower and the actual targeted cells’ photoactivation can be distinguished through *in vivo* imaging, FACS, and *ex vivo* confocal imaging, this presents a complicating factor e.g. when setting gating levels for FACS. Therefore, in this study, we sorted only the brightest population of cells with high levels of mCherry fluorescence (Figures 3B, S7). Future optimization of the phototagging vector and the 2P tagging parameters will help refine photoactivation specificity. Nevertheless, our proof-of-principle implementation of 2P-NucTag to transcriptomically characterize functionally heterogeneous hippocampal PNs that are spatially intermixed in the densely packed pyramidal cell layer demonstrates that 2P-NucTag should be readily applicable to neocortical tissue with lower cell density.

Beyond this first application to the mouse hippocampus, 2P-NucTag will have broad appeal for correlated structural, molecular, and functional analysis of neural circuits. Our approach offers the ability to bridge molecular, cellular-synaptic, and functional architecture in cortical circuits and facilitates the identification of candidate genes that determine the distinct circuit functions of neurons. Our results with ‘Recurring’ place cells further demonstrate that this pipeline should be potent for uncovering molecular signatures of phenomenologically described neuronal subsets with distinct feature selectivity, task-related activity, and longitudinal stability that are spatially intermixed in cortical circuits^21,23,25,99–103^.

Our approach can also be extended in multiple directions. Firstly, as 2P-NucTag is compatible with *post hoc* histological analysis of imaged tissue, it can be combined with spatially resolved transcriptomics^17,18,104,105^ following automated registration between *in vivo* and *post hoc* images^106,107^. Secondly, *ex vivo* electrophysiological recordings from tagged cells will allow for detailed *ex vivo* physiological and anatomical characterization of the *in vivo* tagged cells^9–12^. Thirdly, while we opted to use Meso-seq for our transcriptomic analyses, our pipeline is compatible with sc/snRNA-seq, given the single-cell resolution achieved through targeted photoactivation. Fourthly, further optimization of phototagging with multi-color fluorophores may enable simultaneous labeling of multiple functionally defined cell types in the same animal, while cytosolic tags^48^ can promote anatomical labeling of subcellular axonal or dendritic compartments for downstream structural and connectivity analyses. Lastly, the on-demand nature of photoactivation and its days-long stability following single photoactivation makes our pipeline a valuable tool for precise analysis of transcriptional trajectories following cellular plasticity events during, for example, behavioral learning^53,74,108,109^.

In summary, our novel phototagging approach expands our understanding of the fundamental relationship between the molecular and functional architecture of mammalian cortical circuits and allows for the identification of candidate genes that determine the distinct circuit functions of seemingly homogeneous populations of neurons. Beyond the proof-of-principle implementation to hippocampal circuits, our approach provides a general framework and roadmap for linking genes to cells across neural circuits and model organisms.

## Materials and Methods

### Animals

All animal care and experiment procedures were in accordance with the guidelines of the National Institute of Health. Animal protocols were approved by the Columbia University Institutional Animal Care and Use Committee and the Weizmann Institute of Science Institutional Animal Care and Use Committee. Mice were group-housed under normal lighting conditions in a 12-hour light/dark cycle. *Ad libitum* water was provided until the beginning of training for the spatial navigation task.

### Plasmids and Viral Constructs

pAAV-CW3SL-GCaMP7f-4Ala-H2B-PAmCherry was generated by standard cloning techniques. GCaMP7f was PCR-amplified from Addgene plasmid #104492 with a 3’ Primer that contained sequences encoding four Alanine residues and the P2A sequence (both in frame with the coding sequence of GCaMP7f). H2B-PAmCherry was amplified from Addgene plasmid #133419. The PCR products were then subcloned by Gibson-assembly into Addgene plasmid #61463 after removing EGFP from this plasmid by restriction with ClaI and EcoRI. The sequence of the cloned plasmid was validated by Sanger sequencing and GCaMP7f + 4xAla, P2A and H2B-PAmCherry were all found to be in frame. The plasmid was packaged into AAVDJ at a viral titer of 7.26E+15 essentially as described^55^.

pAAV-hSyn-FLEX-Kir2.2_WT_-T2A-mRuby3 and pAAV-hSyn-FLEX-Kir2.2_Mut-_T2A-mRuby3 were generated at GenScript Inc. by synthesizing the respective inserts (Kir2.2_WT-_T2A-mRuby3, Kir2.2_Mut-_T2A-mRuby3) and cloning them with standard cloning techniques into the viral backbone AAV phSyn1(S)-FLEX-tdTomato-WPRE (Addgene plasmid # 51505). The gene encoding the Kir2.2 channel was synthesized based on Genbank accession number #NM_001267593 (i.e., isoform2 of mouse *Kcnj12*), and mRuby3 based on GenBank accession number #KX987299.1. The plasmid was then packaged into AAVDJ at a viral titer of 1E+13. The strategy of generating active WT channels and Mut control channels were similar to those used in previous studies to generate active and inactive forms of Kir2.1^56,110,111^.

pAAV-EF1a-H2B-PAmCherry was generated by standard cloning techniques. H2B-PAmCherry was amplified from Addgene plasmid #133419 with the relevant sticky ends. The PCR product was then subcloned using Enzymatic ligation into Addgene plasmid #203845 after removing fDIO-mRuby3-2A-dCre from this plasmid by restriction with SalI and EcoRV. The sequence of the cloned plasmid was validated by Sanger sequencing and H2B-PAmCherry were found to be in frame. The plasmid was packaged into AAVDJ at a viral titer of 3.96E+13.

### Surgery

All procedures were performed with mice under anesthesia using isoflurane (4% induction, 1.5% maintenance in 95% oxygen). Mice’s body temperature was maintained using a heating pad both during and after the procedure. Surgeries were performed on a stereotaxic instrument (Kopf Instruments). Before incision, mice were given subcutaneous meloxicam, as well as bupivacaine at the incision site. Doses were calculated based on the animal’s weight. An incision above the skull was made to expose bregma and lambda for vertical alignment. Skull surfaces were cleaned and scored to improve dental cement adhesion. For viral injection, a glass capillary loaded with rAAV is attached to a Nanoject device (Drummond Scientific).

For hippocampal CA1 2P imaging mice except those described in Figure S12 and S13, viruses were injected unilaterally in the left dorsal CA1 at 4 depths using the coordinates: −2.2 AP, −1.75 ML, and −1.2, −1.1, −1.0, −0.9 DV (relative to dura). At each depth, 75nl of AAVDJ-CW3SL-GCaMP7f-4Ala-H2B-PAmCherry was injected. After injection, surgical sites were closed with sutures. Three days after injection, the skull was exposed and a 3mm craniotomy was made centered at the same coordinate of the injection site. Dura was removed, and the cortex was slowly aspirated with continuous irrigation of cold 1X PBS until the fiber tract above the hippocampus was visible. A 3-mm imaging cannula fitted with a 3mm glass coverslip was implanted over the craniotomy site. Cannulas were secured by Vetbond. A custom titanium headpost for head-fixation was secured first with C&B Metabond (Parkell) and then dental acrylic. At the end of each procedure, the mice received a 1.0 ml saline injection subcutaneously and recovered in their home cage with heating applied. Mice were monitored for 3 days after the procedure.

For mice used in Figure S12, 100nl of AAVDJ-hSyn-FLEX-Kir2.2_WT/Mut_-T2A-mRuby3 + AAV9-hSyn-GCaMP8s + AAV1-hSyn-cre were injected at 4 depths using the coordinates: −2.2 AP, −1.75 ML, and −1.2, −1.1, −1.0, −0.9 DV (relative to dura). All other surgery procedures are the same as described above.

For mice used in Figure S13, 100nl of AAVDJ-Ef1a-H2B-PAmCherry + AAV9-hSyn-GCaMP8s were injected at 4 depths using the coordinates: −2.2 AP, −1.75 ML, and −1.2, −1.1, −1.0, −0.9 DV (relative to dura). All other surgery procedures are the same as described above.

For all visual cortex experimental mice, virus was injected unilaterally in the left visual cortex using the coordinates: −2.7 AP, −2.5 ML, and −0.3 DV. 2-3 injections of 300 nl of virus were made using a beveled glass micropipette to target layer 2/3 of the cortex at a rate of 65 nl/min using a microsyringe pump (UMP3T-2, World Prevision Instrument). A 4 mm craniotomy was made above the visual cortex, and a coverglass and custom-made, 3D printed head post painted black to allow for head fixation during imaging was glued to the skull using cyanoacrylate glue (Krazy Glue). Following surgery, animals were administered with analgesic (0.1 mg/kg of buprenorphine and 5 mg/kg of Carprofen). After initial recovery on a heating pad (RWD Life Science), mice were returned to their homecage and monitored for post-op care.

### Behavior paradigm

Mice were first water-deprived and habituated to handling and head fixation at least 7 days after implant surgery. For behavior experiments in Figure 2 and Figure S13, mice were exposed to a 4-m-long linear virtual reality (VR) corridor^65–67^ that stayed consistent in training and recording. For behavior experiments in Figure S12, mice were first exposed to a 4-m VR corridor without features (‘blank’ environment), and they were exposed to the feature-rich VR environment same as the ones used in Figure 2 and Figure S13 on the first day of 2P functional imaging. At the end of the environment, an inter-trial interval of 2 seconds of blank screen was included before the start of the next lap. For the next 10-14 days, mice were trained to run through the virtual environment and lick for a 5% sucrose reward. The rewards were first randomly distributed across the environment, and the number of rewards was slowly reduced from 30 at the beginning of the training to 2 when the mouse was deemed ready for recording. The final reward location was fixed toward the end of the VR environment. Mice were trained to run at least 30-60 laps in the environment. During behavioral imaging, mice were imagined during a single VR session (range: 18-40 min, 26 ± 3 min, n = 9 mice).

### *In vivo* two-photon imaging and data processing

For 2P imaging of dorsal hippocampal CA1, 2P functional imaging was conducted using an 8-kHz resonant scanner (Bruker) and a 16x near-infrared (NIR) water immersion objective (Nikon, 0.8 NA, 3.0-mm working distance). For population imaging in Figure 2 and Figure S12, a field of view of 700 μm x 700 μm was acquired, and for population imaging in Figure S13, a field of view of 420 μm x 420 μm was acquired. All imaging were done at 30 Hz, 512 x 512 pixels using a 940-nm laser (Chameleon Ultra II, Coherent, 45-91 mW after the objective). Red (PAmCherry) and green (GCaMP7f) channels were separated by an emission cube set (green, HQ525/70 m-2p; red, HQ607/45 m-2p; 575dcxr, Chroma Technology), and fluorescence signals were collected with GaAsP photomultiplier tube modules (7422P-40, Hamamatsu). Following the acquisition of two-photon imaging data, Ca^2+^ imaging data was structured and aligned with behavior data using the SIMA analysis package^112^. CA1 ROIs were detected using the Suite2p (v0.14.2) package^113^. To allow detection of all potential ROIs regardless of their activities during the recording, Suite2p was run with Cellpose (‘anatomical_only’)^114^ for the ROI detection step. The pre-trained cyto2 model included in the published Cellpose package was used for ROI detection. When capturing two-photon z-stack images of photoactivated PAmCherry nuclei, a fixed wavelength 1070-nm laser (Fidelity-2W, Coherent) was used for excitation.

For 2P imaging of primary visual cortex, a two-photon microscope equipped with a 12 kHz resonant-galvo scanhead (Bergamo microscope, ThorLabs) was used with a Ti:Sapphire laser (MaiTai DeepSee, Spectra Physics). Imaging data was concatenated, registered, ROIs were drawn and fluorescence was extracted using Suite2p. Following this, all recordings were analyzed using custom software written in house in MATLAB (MathWorks). F signals were neuropil corrected and aligned to timing of visual stimulation using frame time signals collected in Clampex (Molecular Devices) during recording sessions and averaged across repetitions. Field of views (FOVs) were mapped in order to determine the location of V1 and best imaging area for recordings per animal. Receptive fields were mapped by presenting patches of drifting sinusoidal gratings spaced on a 3 by 4 grid of the monitor. Stimuli were presented for 1 s at 0, 100% contrast and 0.04 cpd with an interstimulus interval of 4 s. FOVs were selected based on receptive field mapping and percentage of visually responsive neurons in the area.

### Visual stimulation before and after phototagging

Visual stimuli were generated using the MATLAB Psychophysics toolbox (http://psychtoolbox.org/) and presented on a gamma-corrected LCD screen. The screen was positioned 20 cm from the contralateral eye of the recorded hemisphere. Stimuli were presented for 1 s with an interstimulus interval of 4 s during which a gray screen of mean luminance was presented. Visual stimuli consisted of full-field sinusoidal drifting bar gratings with a temporal frequency of 2 Hz. Varying directions (equal spacing of 30) were presented in a pseudorandom order with 24-25 repetitions per stimulus type.

### *In vivo* two-photon phototagging

Photoactivation was conducted using a three-dimensional random-access acousto-optical (3D-AOD) microscope (3D Atlas, Femtonics)^19,54^. Mice were head-fixed and anesthetized with isoflurane to minimize motion and increase the spatial precision of phototagging. The same 16x NIR water immersion objective was used to find the same field of view as in the functional recordings. Photoactivation was performed at 810-nm (Chameleon Ultra II, Coherent). Two-photon images of the mCherry red fluorescence were taken before and after photoactivation using a 1040-nm excitation laser (Alcor 1040-5W, Spark Lasers). Red (mCherry) and green (GCaMP7f) channels were separated by an emission cube set (green, HQ520/60 m-2p; red, HQ650/160 m-2p; 565dcxr, Chroma Technology), and fluorescence signals were collected with GaAsP photomultiplier tube modules (7422P-40, Hamamatsu). Two-photon images of the mCherry red fluorescence were taken before and after photoactivation using a 1040-nm excitation laser (Alcor 1040-5W, Spark Lasers). For photoactivation, a 7 x 7 μm, 0.1 μm /pixel scanning pattern was placed on the cell to be photoactivated. Each pixel was activated for a total dwell time of 1.3 ms with a laser power of 40 mW. This gave the total scanning time of each cell at 6,370 ms. Following photoactivation, a z-stack was taken for each mouse to assess the photoactivation efficacy.

For phototagging of ‘place cells’ and ‘silent cells’, photoactivation experiments were conducted a day after functional recording sessions. For phototagging of ‘Day 0 cells’ and ‘Recurring cells’, photoactivation experiments were conducted the same day as functional recording sessions. The imaging field of view of 700 μm x 700 μm was matched between the 3D-AOD microscope and the time-averaged GCaMP image from functional recording. After confirming the same field of view as functional imaging, the viewport was zoomed in to a dimension of 250 x 250 μm for more effective identification of targeted cells according to the generated spatial masks. Following photoactivation of all cells, a z-stack image was taken for each mouse to confirm the tagging accuracy. Mice were given *ad libitum* water after functional imaging for at least 12 hours before photoactivation. During the session, mice were monitored every 10 min for breathing rate and reflexes. Heating and eye ointment were applied. Following the photoactivation, mice were returned to the home cage to recover with a heating pad.

For phototagging of neurons in the primary visual cortex, a 405 nm LED light was used to illuminate the field of view for 30 seconds to achieve photoactivation of most of the infected neurons, and photoactivation of neurons were confirmed by recording in the red channel using a 1040 nm laser before and after photoactivation.

### Tissue dissociation and preparation of nuclei for FACS

For all animals used in this manuscript, tissue collection was performed at the same time of the day (5 pm) and at least one hour after the photoactivation. Mice were euthanized using CO_2_. The headpost and metal cannulas were removed. Dorsal CA1 of the hippocampus was collected by first using a 3-mm biopsy punch to cut a circular section of tissue the same size as the craniotomy. A microspatula was used to remove the shallow section of tissue that contained dorsal CA1. The tissue was placed in a 1.5 ml RNase-free Eppendorf tube and snap-frozen in liquid nitrogen. Tissues were stored at −80 ℃ until the start of the nuclei isolation.

Nuclei were prepared for FACS sorting essentially as previously described (*46*). In short, to isolate the nuclei, each collected tissue was transferred to a dounce tissue homogenizer (DWK Life Sciences) with 1ml of homogenization buffer (10 mM Tris Buffer, 250mM Sucrose, 25 mM KCl, 5mM MgCl_2_, 0.1mM DTT, 0.1% Triton X-100, 1X Protease Inhibitor Cocktail, and 40U/μl RNAsin Plus RNase Inhibitor in nuclease-free water). Loose and tight pestles were used to break apart the tissue 10 times each. Following the homogenization, 1 ml of homogenization buffer was added to each dounce, and the homogenate was pipetted up and down to further break apart tissue before being passed through a 30 μm cell strainer (Miltenyi Biotec) and collected in a 15 ml conical tube.

The homogenate was centrifuged at 4 ℃, 700g for 8 min. The supernatants were then removed from the visible nuclei pellet. The nuclei pellet was resuspended with 800 μl of blocking buffer (1X PBS, 1% BSA, and 40 U/μl RNasin Plus RNase Inhibitor in nuclease-free water). The resuspension was incubated on ice for 15 min and transferred to a 1.5 ml tube. 2 μl of CoraLite Plus 488-conjugated NeuN Monoclonal antibody (Proteintech) was added to the resuspension, and incubated on an orbital rotator at 4 ℃ for 30 min. After incubation, nuclei were centrifuged at 4 ℃, 700 g for 8 min. The supernatant was removed, and the nuclei pellet was resuspended with 1000 μl of blocking buffer. DAPI was added to the suspension at a final concentration of 0.001 mg/ml, and the samples were passed through a 40-μm Flowmi Cell Strainer (Bel-Art). All samples were kept on ice until the start of FACS.

### Fluorescence-activated cell sorting (FACS)

The sorting was performed at the Zuckerman Institute Flow Cytometry Core using a MoFlo Astrios Cell Sorter (Beckman Coulter). Event rates were kept between 5000-10000 events per second. The cell sorter uses a linear array of lasers ordered as 640 nm, 488 nm, 561 nm, 532 nm, 405 nm and 355 nm from top to bottom. For experiments described in this manuscript, 488 nm, 561 nm and 405 nm lasers were used to detect the fluorescence of NeuN, mCherry and DAPI respectively. A gating control sample was used to set the gates for DAPI, NeuN and mCherry. Dissociated nuclei were passed through the cell sorter to collect those with high mCherry signals. Bright and Dim mCherry gates were determined after the first 5000 events of DAPI-positive nuclei. Gating was set to collect only the sparse and bright population that showed a high mCherry signal, and these events were collected as the photoactivated nuclei (‘Bright’ mCherry). Nuclei were collected into SMART-Seq CDS sorting buffer that contains 1X lysis buffer, SMART-Seq Oligo-dT and RNase inhibitor. All sorted samples were kept on dry ice as recommended by the SMART-Seq protocol until the start of first-strand synthesis. To increase the likelihood of collecting nuclei in this ultra-sparse population, aborted events for ‘Bright’ mCherry and all events for other positive mCherry were collected into the blocking buffer (200 ml). This suspension was passed through the sorter again with the same fluorescence gating following the completion of the first sorting to capture the bright nuclei.

### Meso-seq

We followed the previously published Meso-seq protocol^55^ with minor modifications. Sorted nuclei were collected in a lysis buffer following the SMART-Seq mRNA LP protocol. Reverse transcription for cDNA was followed by cDNA amplification using 17-18 PCR cycles. Purified cDNA was prepared for sequencing the library using the SMART-seq Library Preparation Kit. Libraries were amplified using 14 PCR cycles. The concentration of the final library was determined using a Qubit3.0 Fluorometer (Invitrogen), and the average DNA fragment size was determined using a Bioanalyzer (Agilent). Sequencing was performed on a NextSeq 2000 sequencer with P2-100 reagents (Illumina). Libraries were diluted and pooled according to the recommendation of the sequencing kit.

### *Ex vivo* electrophysiology in acute hippocampus slices

Mice were transcardially perfused with ice-cold sucrose dissection media (26 mM NaHCO3, 1.25 mM NaH2PO4, 2.5 mM KCl, 10 mM MgSO4, 11 mM glucose, 0.5 mM CaCl2, 234 mM sucrose; 340 mOsm). Brains were then dissected and sliced, while being kept in ice-cold sucrose dissection media, into coronal sections (300 μm thick) containing the hippocampal CA1 using a Leica VT1200S vibratome. Slices were incubated in high osmotic concentrated artificial cerebrospinal fluid (aCSF) (28.08 mM NaHCO3, 1.35 mM NaH2PO4, 132.84 mM NaCl, 3.24 mM KCl, 1.08 mM MgCl2, 11.88 mM glucose, 2.16 mM CaCl2; 320 mOsm) at 32°C for 30 minutes immediately after slicing. Then, slices were incubated in normal osmotic concentrated artificial cerebrospinal fluid (26 mM NaHCO3, 1.25 mM NaH2PO4, 123 mM NaCl, 3 mM KCl, 1 mM MgCl2, 11 mM glucose, 2 mM CaCl2; 300 mOsm) at 32°C for 30 minutes and subsequently at room temperature. All solutions were saturated with 95%-O2/5%-CO2, and slices were used within 6 hours of preparation. Whole-cell patch-clamp recordings were performed in aCSF at 32°C from neurons in the hippocampus. Recording pipettes were pulled from borosilicate glass capillary tubing with filaments (OD 1.50 mm, ID 0.86 mm, length 10 cm) using a P-700 micropipette puller (Sutter Instruments) and yielded tips of 3–5 MΩ resistance. Recordings were sampled at 20 kHz and filtered at 3 kHz. Data were acquired via Clampex10 using a Multiclamp 700B amplifier and digitized with an Axon Digidata 1550B data acquisition board (Axon Instruments). Tagged neurons were identified based on mCherry nuclear fluorescence using a pE-300 white MB LED light (CoolLED) with GYR (525-660nm) spectrum, combined with an Olympus Cy5 Filter Cube Set (ex. 604-644nm; em. 672-712nm).

To ensure that the recorded cells were indeed phototagged neurons, Alexa 594 Hydrazide (10uM) was added to the internal solution to allow co-localization of the fluorescence of the tagged neuron to the one that was patched using confocal imaging in PFA(4%)-fixed slices. For measuring the intrinsic properties, the following internal solution was used: 135 mM k-gluconate, 4mM KCl, 10mM HEPES, 10mM Pcreatine, 4mM Mg-ATP, 4mM GTP-Na and 2mM Na2-ATP. Intrinsic properties were calculated by giving 1.2s long current steps (20pA).

### Tissue collection and processing for *in situ* hybridization

Mice were anesthetized with isoflurane and transcardially perfused with 20 mL of ice-cold 0.01M phosphate base saline (PBS, Sigma) followed by 20 mL ice-cold 4% paraformaldehyde (PFA, Electron Microscopy Sciences) in PBS. Brains were post-fixed in 4% PFA for 24 hours and then saturated with a 10%, 20%, and 30% sucrose solution sequentially over 48 hours until they sunk to the bottom of each successive solution. 30% Sucrose-saturated brains were then embedded in OCT (Optimal Cutting Temperature Compound, Sakura, cat#4583), frozen, stored overnight at −80 ℃, and sliced transversely at 20 µm thickness with a cryostat. Sections were stored at −80 ℃ on slides and used for RNAScope *in situ* hybridization.

### RNAScope Fluorescent In Situ Hybridization (FISH)

20 µm fixed frozen sections of the frozen tissue block were taken and the RNAScope™ Multiplex Fluorescent Reagent Kit v2 – User Manual was followed (cat#: 323100). The *Kcnj12* targeting probe was designed for *Mus musculus* and generated by Advanced Cell Diagnostics Inc (C3: Cat No. 525171, Entrez Gene ID: 16515, GenBank Accession #: NM_010603.6), and same for the *Pcdha8* probe (C1; Cat No. 841011, Entrez Gene ID: 353235, GenBank Accession #: NM_201243.1). Slides were sequentially dehydrated using ethanol solutions of increasing concentrations (50%, 70% and 100%) for 5 min each at RT. 5-8 drops of H2O2 were added to each sample, followed by a 10 min incubation at RT. After rinsing with distilled water, antigen retrieval was performed for 5 min at 99°C (shorten from the manufacturer’s recommendation to preserve PAmCherry fluorescence while maintaining RNAscope signal). To digest sections, RNAscope Protease III was applied to the sections for 30 min at 40°C. The probes were hybridized for 2 h at 40°C and amplified with AMP1 (30 min), AMP2 (30 min), AMP3 (15 min); each was incubated at 40°C. The probe was fluorescently tagged with 1:2000 TSA Vivid Fluorophore 650 (PN 323273). Slides were counterstained with DAPI for a nuclear stain to identify viable cells and mounted in ProLong Gold Antifade Mountant. 20 µm sections were imaged in 3 µm z-steps using an inverted confocal microscope with 20x air objective (A1 HD25, Nikon Instruments Inc.).

## Data analysis

### Quantification and Statistical Analysis

All statistical details for comparisons are described in the text. No statistical methods were used to determine sample sizes. Boxplots show the 25^th^, 50^th^ (median), and 75^th^ quartile ranges with the whiskers extending to 1.5 interquartile ranges below or above the 25^th^ or 75^th^ quartiles, respectively. Outliers are defined as values extending beyond the whisker ranges. For comparisons between two populations with non-normal distributions, the Mann-Whitney U test was used. For comparisons between gene expression datasets, the Wald test followed by multiple corrections via the Benjamini and Hochberg method was used as described in PyDESeq2^115^.

### Event detection

Fluorescence GCaMP traces were deconvolved using OASIS for fast nonnegative deconvolution^116^. As in ref.^117^, these putative spike events were filtered at 3 median absolute deviations (MAD) above the raw trace, using a predetermined signal decay constant of 400 ms. The binarized signal was used to qualify whether a neuron was active at the respective frame. In our analysis, we do not claim to uncover true spiking events in these neurons but use deconvolution for denoising and diminishing Ca^2+^ autocorrelation.

### Spatial tuning curves

The virtual environment was divided into 100 evenly spaced bins (4 cm), which were then utilized to bin a histogram of each cell’s neuronal activity. Neuronal activity was filtered to include activity from when the animal was running above 3 cm/s and to exclude activity during the 2-sec teleportation at the end of the 4-m track. The spatial tuning curves were normalized for the animal’s occupancy and then smoothed with a Gaussian kernel (σ = 12 cm) to obtain a smoothed activity estimate.

### Place cell detection

Place fields were detected by identifying locations in the virtual environment where a neuron was more active than expected by chance. We circularly shifted each neuron’s deconvolved spike trace and recomputed the smoothed, trial-averaged spatial tuning curve of the shifted trace to generate a shuffled null tuning curve per cell. We repeated this procedure 1000 times in order to calculate the 95^th^ percentile of null tuning values at every spatial bin to generate a threshold for a p<0.05 significance curve. Spatial tuning curves that surpassed the null threshold were marked as candidate place fields, and the place field width was calculated as the total bins where the tuning curve exceeded the shuffled null tuning curve. To restrict our analysis to neurons with specific firing fields, we additionally required that place fields have a width greater than 8 cm and less than one-third of the virtual environment (1.3 m). To ensure that the place field activity was stable, we also required that all place cells had activity for at least 20 laps.

If the binarized signal trace for a cell did not have any detected events via OASIS deconvolution, the cell was classified as silent.

For experiments described in Figure S12, five consecutive days of recording were first concatenated as one image sequence and Suite2P was used for motion correction and ROI-detection. They were then separated into recording for each day for analysis. Place fields detection were performed for each cell for each recording days, and same ROI number were used across days. Cells with place field for at least 4 out of 5 days were classified as ‘Recurring’ cells. ‘Day 0’ cells were place cells with place fields on the first recording day, done in a separate set of animals.

### PAmCherry fluorescence quantification

(Figure 1C,D)

Red PAmCherry fluorescence was calculated as the tagged nuclei fluorescence versus the background fluorescence of nearby untagged nuclei for the image taken post-tagging. This was intentional to control for periodic two-photon imaging at 940 nm that may cause an increase in fluorescence for every cell in the FOV.

### Fluorescence intensity distribution analysis

(Figure 1F, right)

Red PAmCherry tagged nuclei were segmented using Cellpose with manual curation performed within the Cellpose GUI for the max axial projections of the *ex vivo* z-stack (excitation: 568 nm nm) and *in vivo* z-stack (1070 nm) to generate masks. The masks were used to segment the 3-D volumes of each cell and the respective fluorescence profiles were normalized and aligned based on the peak value. The average and standard error were computed based on the aligned fluorescence profiles of the cells from the respective *in vivo* and *ex vivo* volumes.

### Two-photon background excitation fluorescence quantification

[Figure 2G(2P imaging), Figure S4F (2P imaging)]

Changes in PAmCherry red fluorescence due to two-photon fluorescence excitation at 940 nm from functional recordings were examined. By taking the average fluorescence of the pre-imaging red channel image and post-imaging red channel image detected at 1070 nm, ΔF/F was computed using the change in fluorescence between the average post-imaging red image and pre-imaging red image divided by the pre-imaging red image average [(post – pre) / pre].

### Tagged and Off-target Fluorescence Change Quantification

[Figure 1G (target, off-target), Figure S4F (target, off-target)]

Tagged nuclei were segmented using Cellpose with manual curation performed within the Cellpose GUI for the max axial projection of the *in vivo*-stacks from each mouse to generate masks. These masks were used to segment a 3D volume for each nucleus that was targeted across all mice. The number of off-target nuclei was manually quantified laterally and axially per targeted nuclei. Masks for off-target nuclei were hand-drawn on the max axial projection to exclude the targeted nuclei and were at most 2 nuclei bodies away. Experimental background fluorescence masks were drawn on the surrounding areas with successfully tagged nuclei and excluded all targeted and off-target nuclei.

The fluorescence values for all tagged and off-target nuclei were percentile-filtered to exclude the lower 10% of fluorescence values to account for vignetting effects and mask inhomogeneities over the max axial projection. Background fluorescence levels were selected in the areas of field of view without tagged nuclei. The percentile filtered fluorescence values for tagged and off-target nuclei were averaged, and the ΔF/F was computed by taking the difference between the average tagged or off-target fluorescence value and the average background fluorescence and then normalizing by the average background fluorescence [tagged: (tagged – background)/background; off-target: (off-target – background)/ background]. Note this background value is distinct from the one described in the two-photon background excitation fluorescence quantification.

### GCaMP quantifications

[Figure S1AB, Figure S4I]

Frequency: The total number of deconvolved events by OASIS was normalized by the total duration of the recording.

For each cell, the average transient was segmented within a 15-second time window and computed by averaging along aligned deconvolved spike times. If multiple detected events were within 50 frames, the events were treated as a single transient. A cell was only used if there were at least 3 detected transients within the total trace to exclude cells without obvious GCaMP-Ca^2+^ dynamics. For each average transient, we computed the median value based on 5 seconds pre peak transient and the range 5-10 seconds after the peak transient (given that the average transient took significantly less than 5 seconds to resolve) to act as the baseline.

Amplitude: The difference between the max value of the average transient and the baseline value was computed Half-Rise Time: The time between the half-max value prior to the max and the max value of each transient was computed. We excluded cells where the average half-max value was not observed prior to the transient and performed 99^th^ percentile filtering to remove extreme outliers. Due to sampling rate limitations, we cannot comment on the true half-rise time, so these are approximations.

Half-Decay Time: We computed the time between the max and the half-max value following the transient peak. To remove extreme outliers, we performed 99^th^ percentile filtering.

### Visually evoked neural activity

[Fig. S6]

Visually evoked responses were calculated as the mean ΔF/F during visual stimulus presentation:

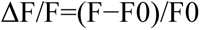

Where F0 was calculated as the mean fluorescence during the 1 second preceding the visual stimulus. To determine if a cell was visually responsive, a one-way non-parametric ANOVA (Kruskall-Wallis) was computed comparing the evoked responses to baseline fluorescence. An additional non-parametric test was performed between the activity during the interstimulus interval and during presentation of the preferred visual stimulus (Wilcoxon Rank Sum test, p < 0.05).

Neurons were included in further analyses if they were visually responsive both before and after photoactivation, and if they were determined to be successfully photoactivated.

Orientation selectivity was calculated using a calculation of the global orientation selectivity index (gOSI):

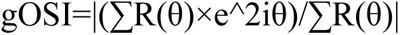

Where R(qk) is the response to angle qk. To calculate the population average, responses of each cell were normalized to their maximum response and artificially centered to 0. Normalized responses of cells were averaged and fitted using a double Gaussian:

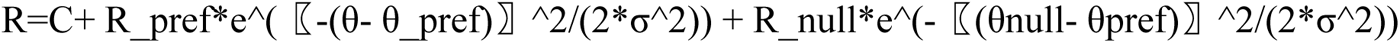

Where C is a constant offset, Rpref is the response to the stimulus that produced the maximum response and Rnull is the mean response to the opposite direction: Orientation preference was calculated as the orientation that elicited the highest dF/F value according to the fit of the gaussian tuning curves.

### Image denoising

To correct for vertical scanning artifacts, we utilized combined wavelet and Fourier filters described in ref.^118^ [github: https://github.com/DHI-GRAS/rmstripes]. Symlet 20 wavelets were used with varying levels of decomposition (2-5) for discrete wavelet transform to perform vertical striping correction in static images.

### *In vivo* and *ex vivo* image registration

The *in vivo* and *ex vivo* images were transformed into 3D volumetric images for registration. The *in vivo* sequential images were concatenated across the z direction, i.e., depth, stacking the 2D images into a volumetric representation using MATLAB. The *ex vivo* images composed of 2D slices in each section were concatenated into 3D volumetric data using a stitching algorithm developed as a precursor for automatic *ex vivo* and *in vivo* registration^106^

[github: https://github.com/ShuonanChen/multimodal_image_registration]. The discontinuity between *ex vivo* sections result in an unknown spatial correlation between them, requiring registration between sections. Common cells between sections (i.e., sections one and two) were used as reference markers for registration. The common cells were manually selected using a Napari GUI in Python and were utilized as reference markers to inform the scaling and affine transformations to be applied. The scaling and affine transformations were run automatically, transforming the second section to align with the first section. In the scaling transformation, the relative distances between cells in a section were compared to the cells in the first section, inducing an enlargement or shrinkage of the second section to match the first. In the affine transformation, the second section was geometrically transformed to align correctly with the first section. The transformations were obtained and applied to each slice in each section, and then each slice was concatenated together across the z direction to form a volumetric image.

The *in vivo* and *ex vivo* 3D images were then adjusted to have a uniform pixel size in all dimensions (1 μm in x,y and z), ensuring matching FOVs, and equivalent resolution across both images. Time-averaged representations of *in vivo* and *ex vivo* volumetric stacks were attained by employing maximum intensity projection (MIP) representations in FIJI, compressing the stacks into 2D images^119^. The *in vivo* and *ex vivo* registration was carried out using a non-rigid registration algorithm for the (i) 3D volumetric stacks and (ii) MIP (2D) images. All cells common to both *in vivo* and *ex vivo* images were manually selected as the centroid of each cell using a Napari GUI in Python. The common cells were utilized as features to inform the scaling, affine transformation, and deformation transformation, which were applied to the *ex vivo* image. The scaling and affine transformations were run automatically. In the scaling transformation, the distances between the cells in the *ex vivo* image and matching cells in *in vivo* induce enlargement or shrinking of the *ex vivo* image to match the *in vivo* image. In the affine transformation, the *ex vivo* image was geometrically transformed to align with the *in vivo* image using the matching cells. The images in the GUI are updated to reflect the changes induced by the scaling and affine transformation. The deformation transformation uses a vector field, smoothed with Gaussian filtering, to move cells and deform the image, ensuring features in the transformed *ex vivo* image align with the *in vivo* image. The deformation transformation was iteratively employed, beginning with a Gaussian kernel size of 100, reducing to a kernel size of less than 10, with the user inspecting the alignment and making manual adjustments to the cell centroid position in the GUI. The completed transformed *ex vivo* image was then overlaid with the *in vivo* image.

### RNA-sequencing data analysis

Sequencing data from the Illumina Sequencer was first post-processed through the Illumina DRAGEN secondary analysis pipeline to de-multiplex based on a unique index for each sample. RNA-seq reads were aligned to the mouse genome (mm39) using STAR^120^. Unique reads were counted using HTSeq^121^. Gene counts were then normalized with trimmed mean of M values (TMM) normalization. HTSeq generated reads were then analyzed for differential expression using PyDESeq2^115^. Following HTSeq counts, any genes with expression in less than 3 samples were discarded. FDR adjusted p values were used to determine significantly different genes. Both ‘place’ and ‘silent’ cell samples were analyzed against randomly tagged, function-blind, and sex-matched samples generated from mouse CA1 tissue in the same way as described in this methods section above. The same comparisons were made to identify differentially expressed genes for ‘place’ versus ‘random’, and ‘silent’ versus ‘random’. Within this ‘random’ dataset, male-female samples were compared to identify differentially expressed genes influenced by sex. All DEGs from these comparisons were cross-referenced to find common hits.

### Gene set enrichment analysis

Gene set enrichment analysis was performed on HTSeq-generated counts using the gseapy package^122^ in Python 3.11. Enrichment of the fifty hallmark pathways from the molecular signatures database for *Mus musculus* (version 2023.2) was compared in place and silent cells. Comparisons were done with t-test and 1000 permutations.

### RNAscope FISH signal quantification and statistical analysis

Cell detection was performed using Cellpose, which identified individual cells based on DAPI nuclear staining. This segmentation was manually curated. The detection and quantification of RNAscope probe signals was performed using QuPath’s Subcellular Detection tool (version 0.5.1). The detection threshold was set to ensure accurate identification of signal dots. To compare mean RNA expression levels between tagged and non-tagged cells, a Mann-Whitney U Test was used. For non-tagged control cells, a random subset was selected from the same tissue sections and CA1sublayer as the tagged cells, with the number of control cells matched to the number of tagged cells for each animal.

## Acknowledgments

We thank Ira Schieren and Max Wallach (Columbia, Zuckerman Institute, ZI) and Dr. Efrat Hagai (Flow Cytometry Unit, Weizmann Institute of Science) for help with FACS, Columbia University Herbert Irvine Comprehensive Cancer Center Molecular Pathology Shared Resources for help with the Bioanalyzer, the Bendesky lab in ZI and Dr. Hadas Keren-Shaul, Revital Ronen at the Weizmann, Nancy and Stephen Grand Israel National Center (G-INCPM) for help with RNA-sequencing, George Zakka (Losonczy Lab) for technical support, Erica Rodriguez (Salzman lab) in ZI for help with tissue processing, the Lomvardas lab in ZI for sequencing advice and help, the Gogos lab in ZI for molecular bench space, the Polleux lab in ZI for help with cell counter and confocal microscopy. We thank Franck Polleux, Joseph Gogos, Steven A. Siegelbaum, and Darcy Peterka (Columbia, ZI) for feedback on the manuscript.

## Author contributions

Conceptualization: JS, BN, DK, AL, IS

Methodology: JS, BN, DK, BR, SAH, ET, KCK, MECP, HCY, BMS, AX

Investigation: JS, BR

Software: BR, JS, TSM, CKO, EV

Formal analysis: JS, BR, TSM, ET, AL

Validation: JS, BN, DK, KCK, BR

Visualization: JS, BR, TSM, SAH, CKO, ET, MECP, KC-KM

Data curation: JS, BR, ET, TSM

Funding acquisition: AL, IS

Resources: AL, IS

Project administration: AL, IS

Supervision: AL, IS

Writing – original draft: JS, BN, BR, TSM, AL, IS

Writing – review & editing: all authors

## Data and material availability

All data, code, and materials used to generate figures and perform statistical tests are available at Zenodo or NWB upon publication. RNA-seq data will be made freely available on NCBI GEO upon acceptance of our manuscript. Accession numbers for publicly available reagents are included in supplementary materials and methods.

## Supplementary Figures

**Figure S1.**
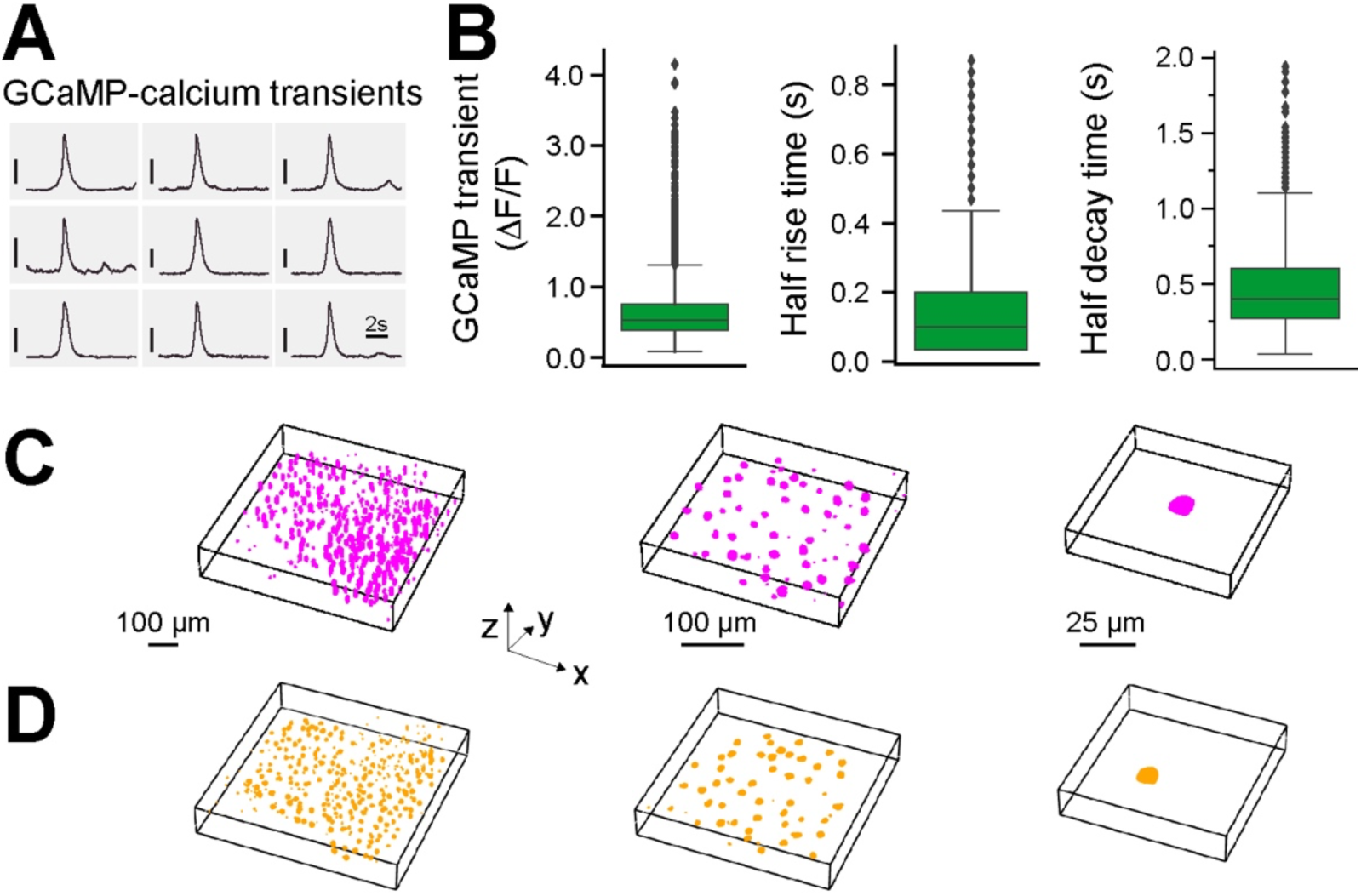
Additional data on 2P-NucTag. (**A**) Representative average GCaMP-Ca^2+^ transients from nine CA1 PNs, vertical scale bar (50% ΔF/F). (**B**) Left: GCaMP amplitude (54.6% ± 0.3% ΔF/F, n = 8190 cells in 9 mice). Middle: GCaMP half rise time (0.15s ± 0.001s, n = 7797 cells in 9 mice), Right: GCaMP half decay time (0.44s ± 0.003s, n = 8282 cells in 9 mice, 940 nm excitation, See methods for cell exclusion criterion). Boxplots show the 25^th^, 50^th^ (median), and 75^th^ quartile ranges, with the whiskers extending to 1.5 interquartile ranges below or above the 25^th^ or 75^th^ quartiles, respectively. Outliers are defined as values extending beyond the whisker ranges. (**C**) Left: *in vivo* 3D visualization of the entire field of view (FOV). Middle: subset of *in vivo* 3D visualization. Right: representative cell from *in vivo* 3D visualization. (**D**) Left: *ex vivo* confocal 3D visualization of entire FOV. Middle: subset of *ex vivo* 3D visualization. Right: representative cell from *ex vivo* 3D visualization. *C* and *D* are corresponding to Figure 1F.

**Figure S2.**
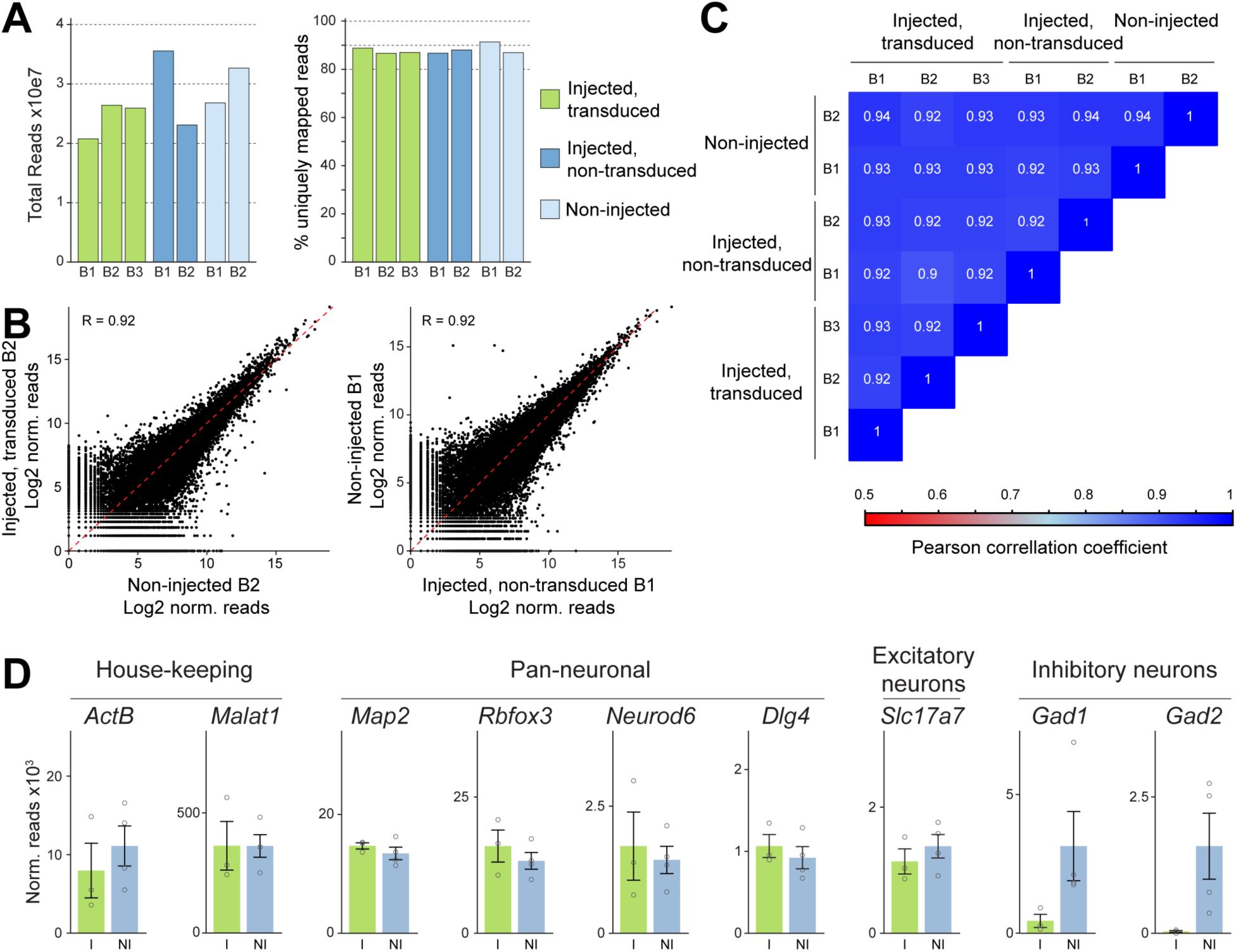
Meso-seq on hippocampal CA1 pyramidal neurons with 2P-NucTag construct. (**A**) Total reads and percentage of uniquely mapped reads from all libraries (B = biological replicate). (**B,C**) Pearson-correlations of all pair-wise comparisons of the expression levels of all expressed genes in the 7 libraries. (**B**) Two example comparisons. (**C**) Correlation coefficients of all comparisons (B = biological replicate). (**D**) Expression values of example genes in non-transduced neurons (NI = pool of “Non-injected” and “Injected, non-tranduced”) and transduced neurons (I = “Injected, transduced”) (error-bars = SEM).

**Figure S3.**
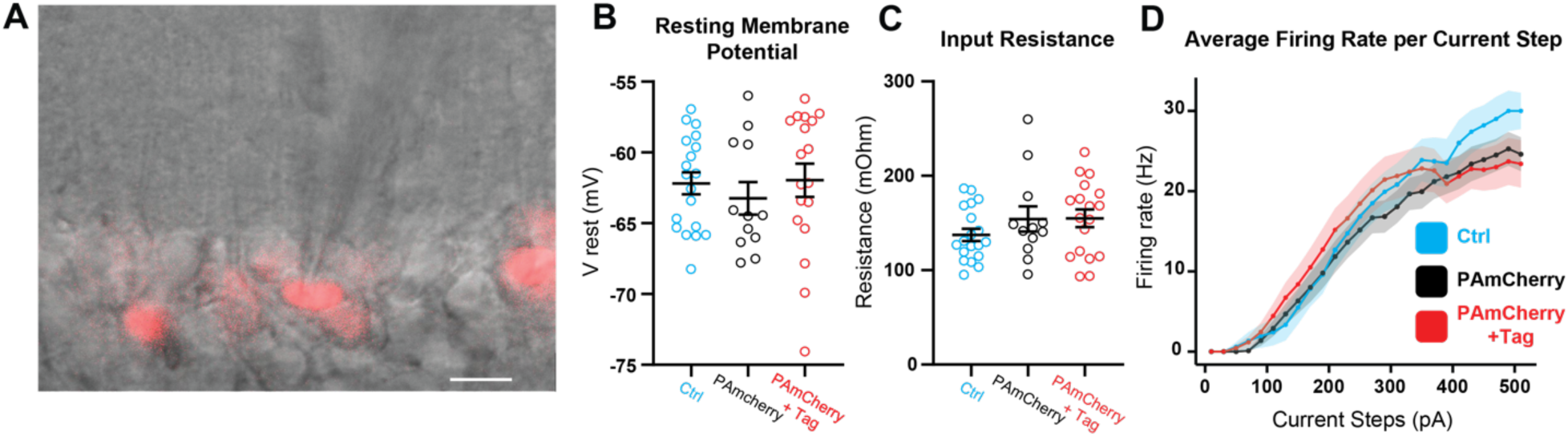
The 2P-NucTag construct does not alter the intrinsic properties of the CA1 pyramidal neurons. (**A**) Representative image of CA1 pyramidal neurons with (red fluorescence) and without photoactivation during whole cell patch clamp recording. Scale bar: 10 μm. (**B,C**) Quantified data of Resting membrane potential and Input resistance in CA1 excitatory neurons. Blue - control neurons (*Ctrl*), recorded in CA1 of the contralateral non-infected hemisphere. Black- are the neurons recorded from the infected hemisphere but have not been phototagged (*PAmCherry*). Red (*PAmCherry+Tag*) are the phototagged neurons (Ctrl n = 19 cells from 7 mice; PAmCherry n = 12 cells from 8 mice; PAmCherry+Tag n = 18 cells from 7 mice; Statistics Kruskal-Wallis test with post-hoc Dunnett’s T3 multiple comparisons test: no statistically significant differences were observed). (**D**) Average firing rate per current step (F-I curve) in each condition. (error bars represent SEM in all data panels).

**Figure S4.**
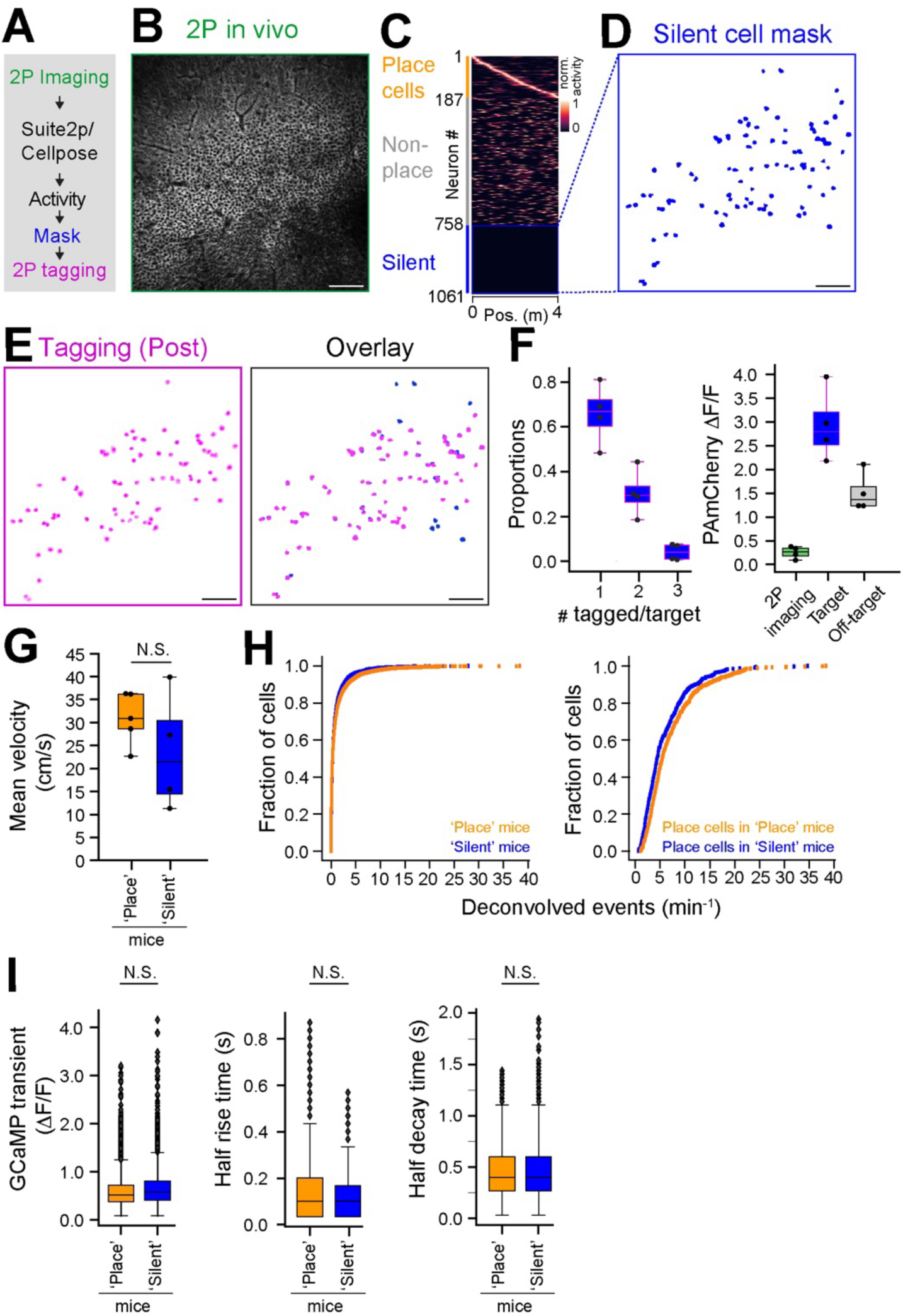
*In vivo* 2P-NucTag of silent cells. (**A**) Pipeline for two-photon (*2P*) phototagging of functionally identified ‘silent’ neurons in CA1 during VR spatial navigation (as in Figure 1B). (**B**) Example 2P imaging field of view (FOV) of GCaMP in the CA1 pyramidal layer from a mouse with “silent” cells targeted. Scale bar: 100 μm. (**C**) CA1PNs detected with Suite2p/Cellpose in the FOV shown in *B*. (**D**) Spatial mask (blue) of identified ‘silent’ cells from the FOV in *B.* Scale bar: 100 μm. (**E**) Left: PAmCherry fluorescence (magenta) of tagged nuclei after 2P phototagging. Right: overlay of spatial masks of identified CA1PNs and tagged nuclei for the FOVs in *D* and *E.* Note that we only tagged a subset of silent cells present in the FOV. Scale bar: 100 μm. (**F**) Left: Proportion of single, double, and triple-tagged nuclei following phototagging of a single silent cell. Right: relative PAmCherry fluorescence change for non-tagged cells in the FOV after 2P imaging (green), after 2P phototagging of targeted silent cell nuclei (blue) and off-target nuclei (gray, n = 4 mice). (**G**) Average velocity of the mice during virtual reality navigation task (Mann-Whitney U test, p-value = 0.286). (**H**) Left: deconvolved events per minute from all cells across all mice from 2P GCaMP-Ca^2+^imaging (averaged across mice, n = 5 ‘Place’ mice, n = 4 ‘Silent’ mice, Mann-Whitney U Test, p-value = 0.28). Right: deconvolved events per minute from place cells across all mice (averaged across mice, place n = 5, silent n = 4, Mann-Whitney U Test, p-value = 0.90). (**I**) Left: GCaMP transient amplitude of all cells between groups (averaged across mice, n = 5 ‘Place’ mice, n = 4 ‘Silent’ mice, Mann-Whitney U Test, p-value = 0.14), Middle: GCaMP half rise time of all cells between groups (averaged across mice, n = 5 ‘Place’ mice, n = 4 ‘Silent’ mice, Mann-Whitney U Test, p-value = 0.14), Right: GCaMP half decay time of all cells between groups (averaged across mice, n = 5 ‘Place’ mice, n = 4 ‘Silent’ mice, Mann-Whitney U Test, p-value = 0.81). Boxplots show the 25^th^, 50^th^ (median), and 75^th^ quartile ranges, with the whiskers extending to 1.5 interquartile ranges below or above the 25^th^ or 75^th^ quartiles, respectively. Outliers are defined as values extending beyond the whisker ranges.

**Figure S5.**
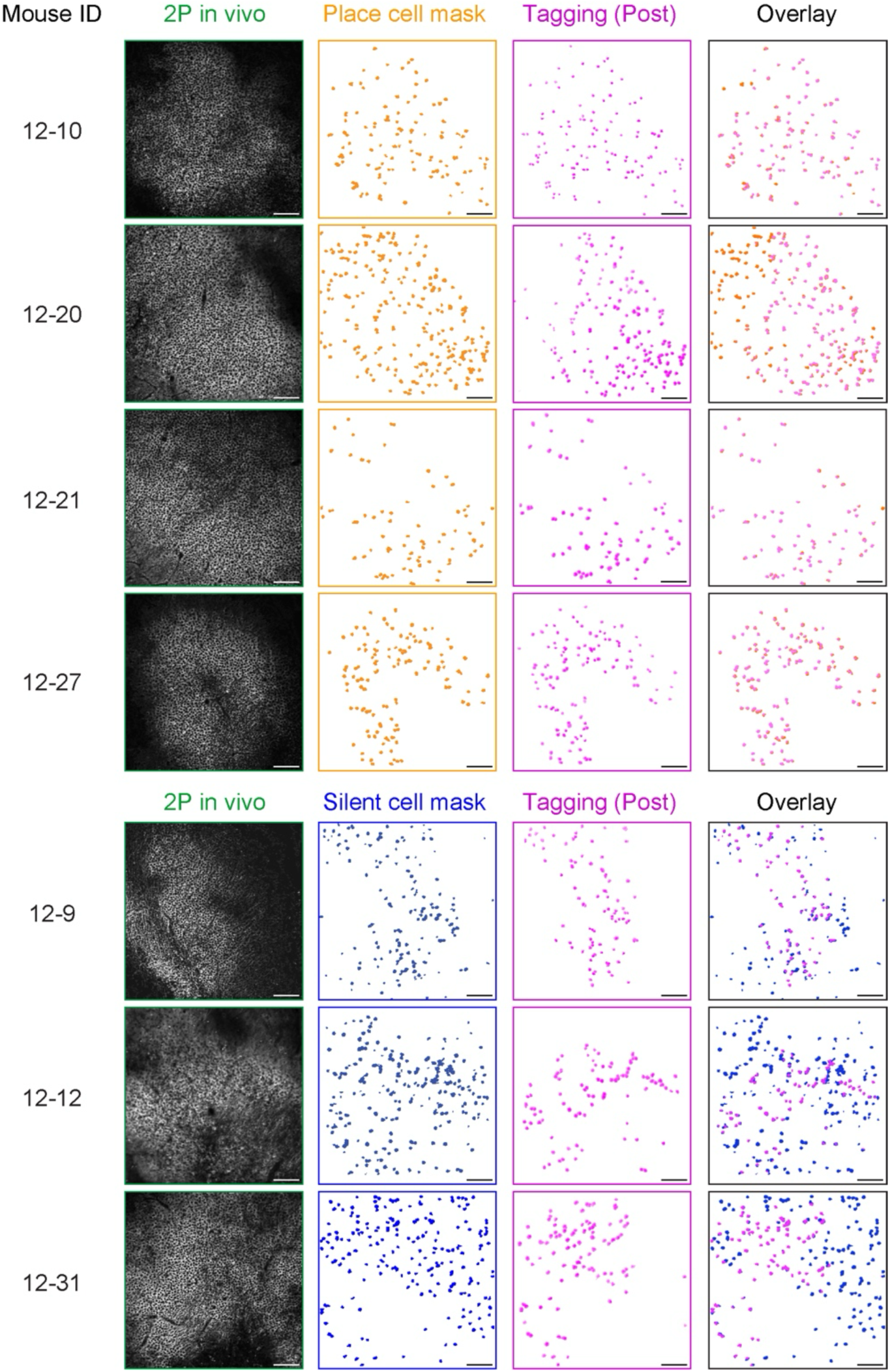
Additional data on phototagging. From left to right by column: animal ID (animals in Figure 2 and *Figure S2* are not shown here), *in vivo* two-photon (*2P*) imaging fields of view (FOVs), functionally defined masks (orange for place cells and blue for silent cells), PAmCherry fluorescence (magenta) of tagged nuclei after 2P phototagging and overlay of spatial masks from identified CA1PNs and tagged nuclei for the respective FOV in the same row. Note that we only tagged a subset of silent cells present in the FOV in ‘Silent’ mice, in order to approximate the number of phototagged place cells in ‘Place’ mice. Scale bar: 100 μm.

**Fig. S6.**
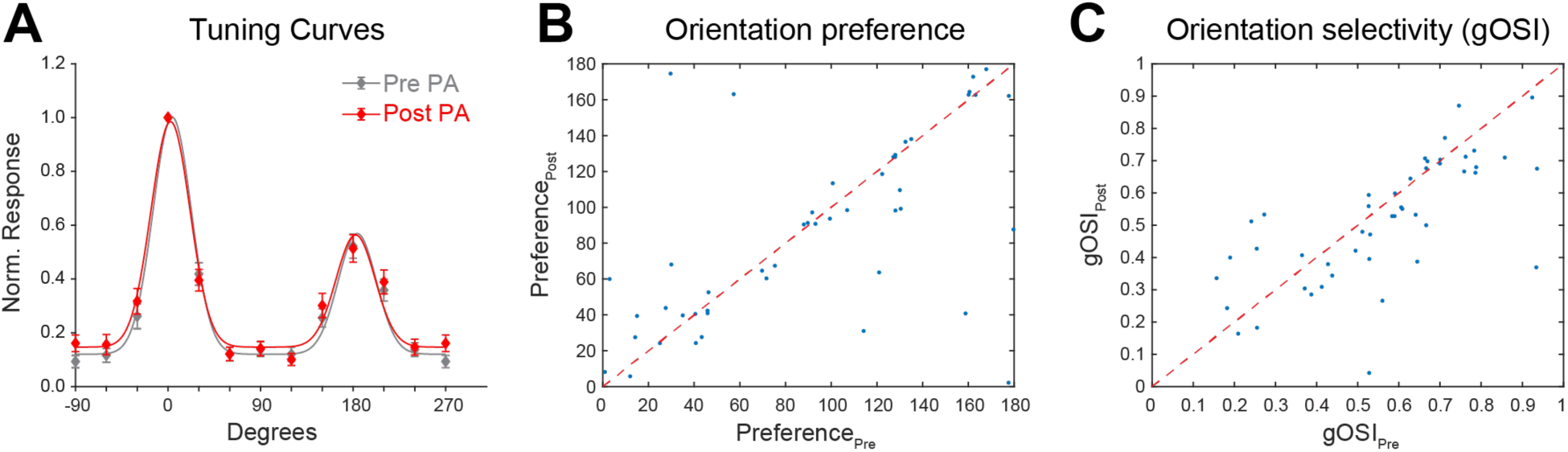
Orientation tuning of visually responsive neurons in V1 before and after photoactivation. (**A**) tuning curve for visually responsive neurons before and after photoactivation (Gray – before photoactivation, Red – after photoactivation). Line - normalized double-gaussian-fitted population averages (**B**) Orientation preference visually responsive neurons before and after photoactivation. (**C**) Orientation selectivity before and after photoactivation. (Error bars represent SEM in all data panels).

**Figure S7.**
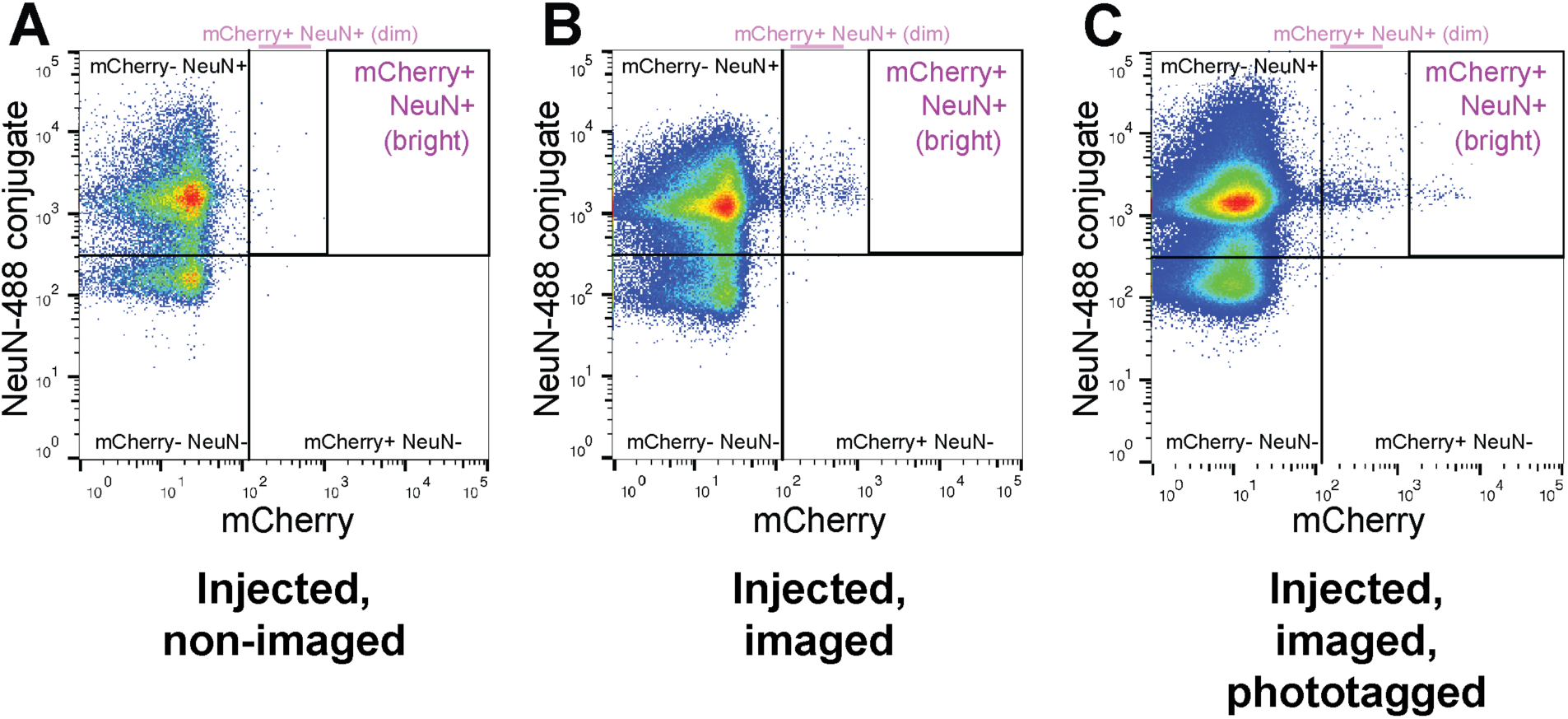
FACS Gating based on mCherry fluorescence. (**A**) representative FACS graph for samples that were injected but not imaged. (**B**) samples that were injected and imaged. (**C**) samples that were injected, imaged, and phototagged.

**Figure S8.**
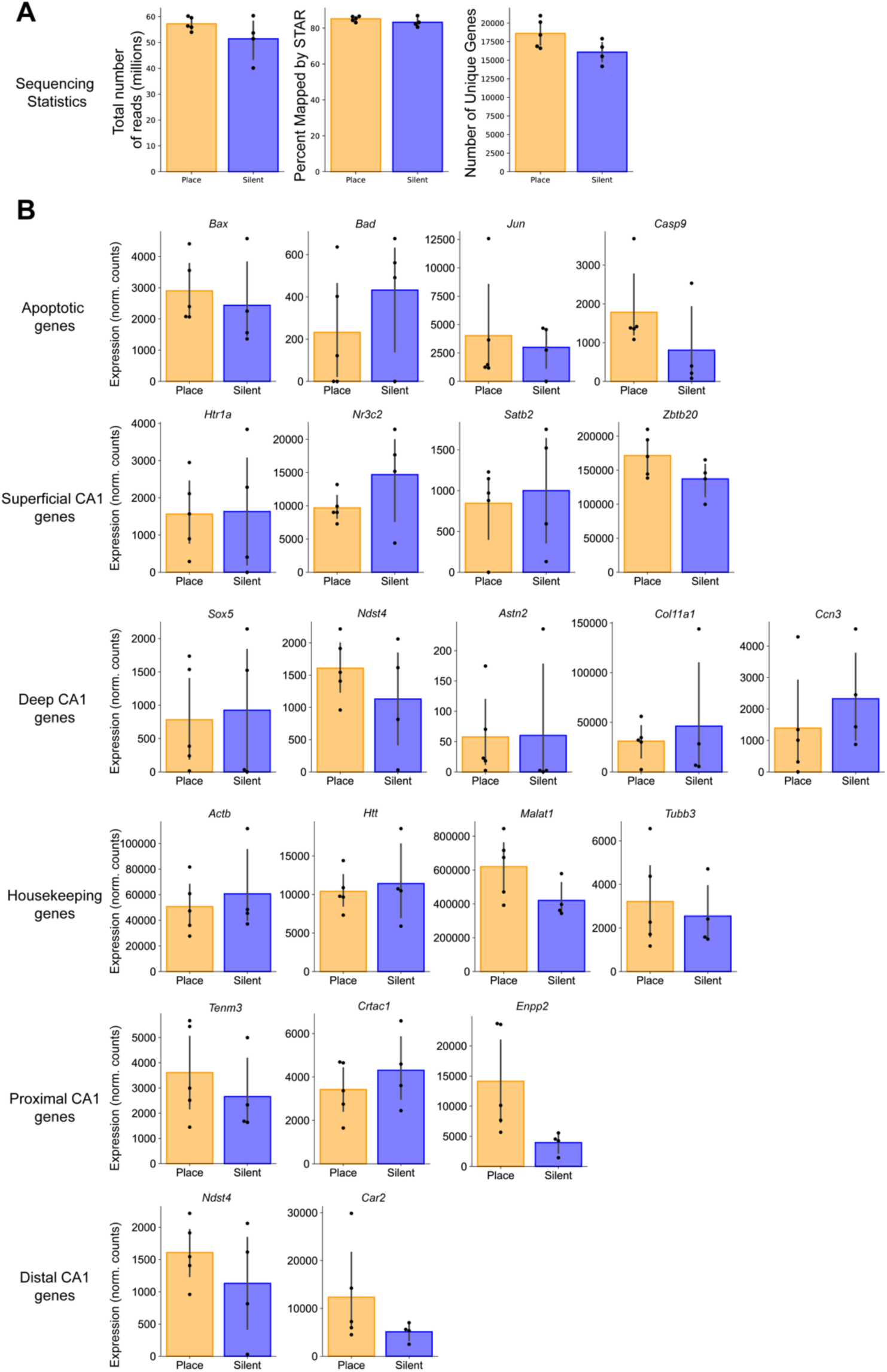
Additional data on transcriptomics analysis of place and silent cells. (**A**) Sequencing statistics for all ‘place’ and ‘silent’ cell samples. Total number of reads - 40 to 60 million reads, 54.64 ± 2.09, n = 9. Percent mapped by STAR - 80.58 to 86.92%. 84.33 ± 0.73. Number of unique genes - 14176 to 20990, 17486 ± 714. (**B**) Normalized counts for groups of genes plotted for ‘place’ versus ‘silent’. Here we show that gene expression of apoptotic genes, superficial CA1 genes, deep CA1 genes, housekeeping genes, proximal CA1 genes, and distal CA1 genes are not different between the two groups (FDR adjusted p-value. *<0.05, \*\***<**0.001, ***<0.001, PyDeSeq2. All comparisons in this figure are not significant).

**Figure S9.**
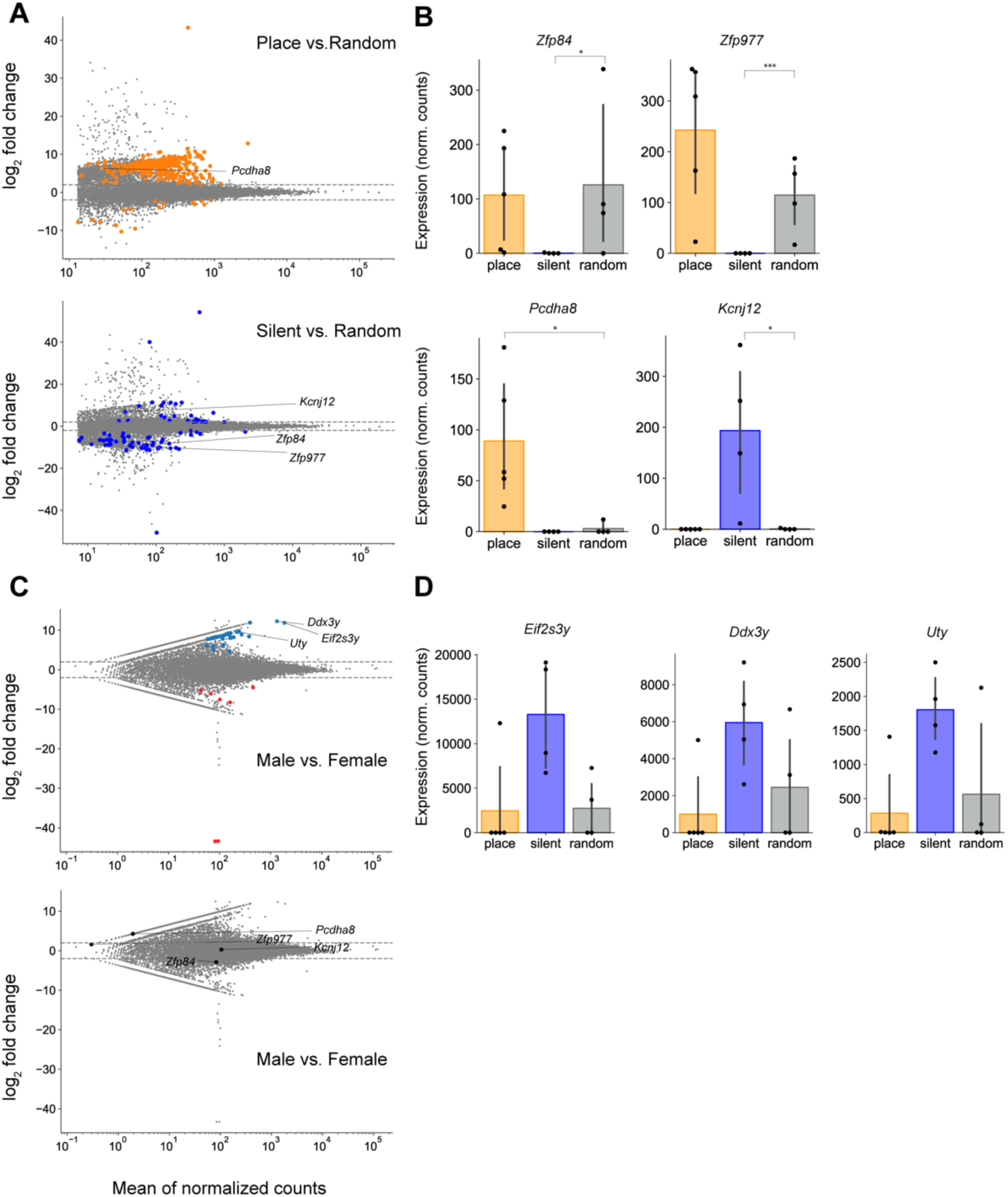
Additional data on transcriptomics analysis of place and silent cells. (**A**) Top: MA plot of ‘place’ versus ‘random’. Differentially expressed genes (DEGs) are labeled in orange. Bottom: MA plot of ‘silent’ versus ‘random’. DEGs are labeled in blue. Both: DEGs that are common for ‘place’ versus ‘random’ were highlighted and labeled. (**B**) Normalized counts for 4 example genes that are significantly differentially expressed across comparisons (FDR adjusted p-value. *<0.05, \*\***<**0.001, ***<0.001, PyDeSeq2. Showing here a comparison of ‘place’ versus ‘random’ or ‘silent’ versus ‘random’. ‘Place’ versus ‘silent’ comparisons were shown in Figure 3). (**C**) MA plot of male versus female for the ‘random’ dataset. Top: Y-linked genes that are differentially expressed between sex are highlighted and labeled. Bottom: same four genes in panel A and B are highlighted and labeled. They are not differentially expressed between sex (FDR adjusted p-value. *<0.05, \*\***<**0.001, ***<0.001, PyDeSeq2. Otherwise, comparisons are not significant). (**D**) Normalized counts for 3 example DEGs between male and female.

**Figure S10.**
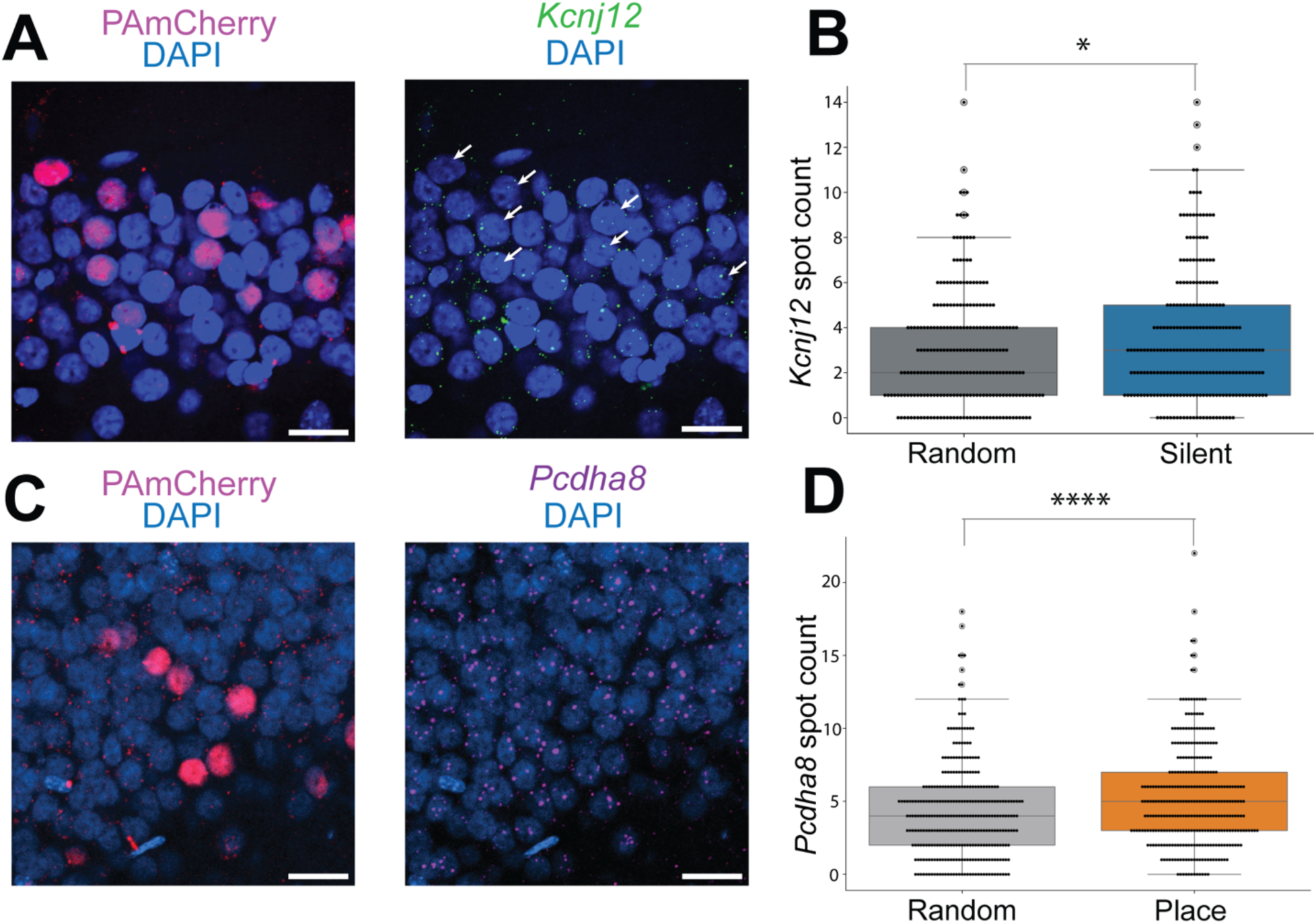
Spatial distribution of selected gene transcripts. (**A**) Left: Confocal horizontal image showing tagged silent cells expressing PAmCherry (red with nuclei counterstained by DAPI (blue). Scale bar: 20 μm. Right: Confocal image of the same tagged tissues as in (A), hybridized with a probe for *Kcnj12* transcripts (green) using RNAscope Multiplex Assay v2. Nuclei are counterstained with DAPI (blue). Scale bar: 20 μm. For images shown in A&B, red mCherry image was obtained at 20x zoom pre-RNAScope; blue DAPI channel and green Kcnj12 channels were obtained at 20x zoom post RNAScope. (**B**) Box plot comparing the expression levels of *Kcnj12* in tagged cells (n=247, n=3 mice, median=3, IQR=4) and randomly selected non-tagged cells (n=247, n=3 mice, median=2, IQR=3). Mann-Whitney U Test, p-value = 3.16e-2). **(C)** Left: Confocal horizontal image showing tagged place cells expressing PAmCherry (red with nuclei counterstained by DAPI (blue). Scale bar: 20 μm. Right: Confocal image of the same tagged tissues as in (A), hybridized with a probe for Pcdha8 transcripts (magenta) using RNAscope Multiplex Assay v2. Nuclei are counterstained with DAPI (blue). Scale bar: 20 μm. (D) Box plot comparing the expression levels of *Pcdha8* in tagged cells (n=321, n=3 mice, median=5, IQR=4) and randomly selected non-tagged cells (n=321, n=3 mice, median=4, IQR=4). Mann-Whitney U Test, p-value = 1.58e-4). Boxplots show the 25th, 50th (median), and 75th quartile ranges, with the whiskers extending to 1.5 interquartile ranges below or above the 25th or 75th quartiles, respectively. Outliers are defined as values extending beyond the whisker ranges.

**Figure S11.**
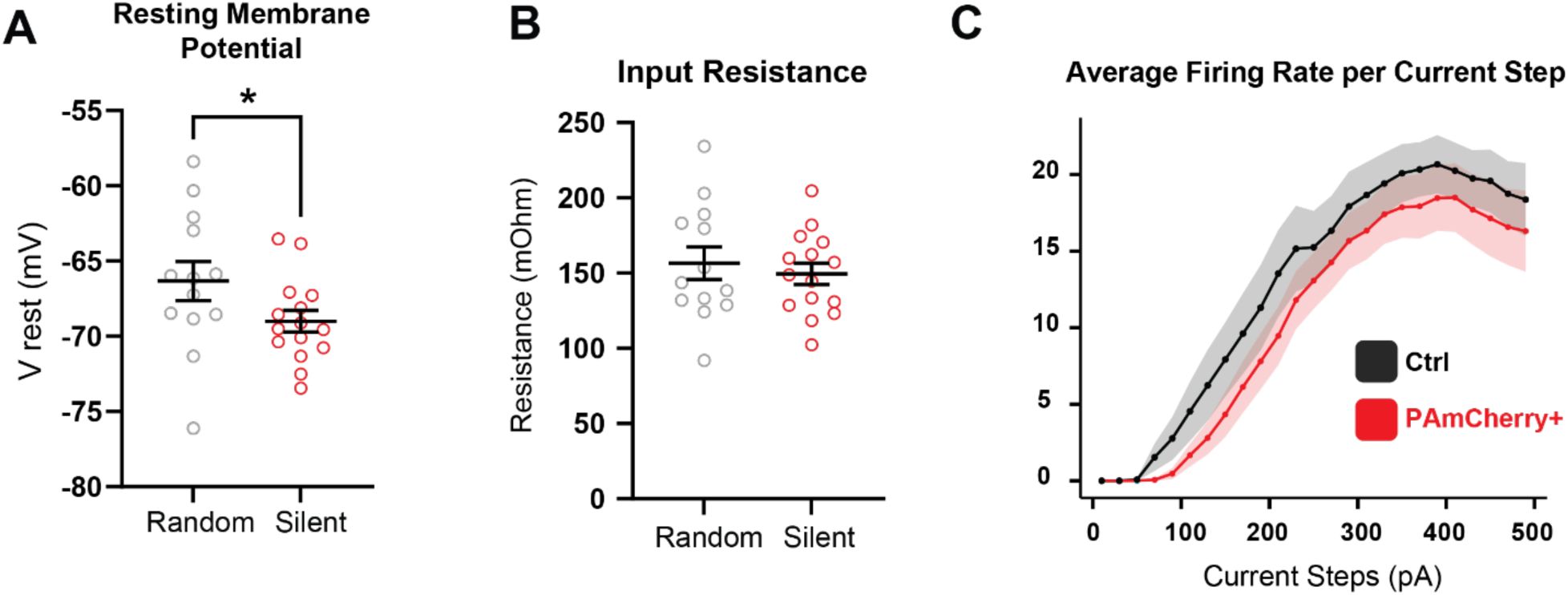
Silent cells display a decrease in resting membrane potential. (**A,B**) Quantified data of Resting membrane potential and Input resistance in Random (non-silent, *Ctrl*) and silent (PAmCherry+) CA1 PNs (Random n = 19 cells from 4 mice; Silent n = 12 cells from 4 mice). (Mann-Whitney U test. Resting Membrane Potential: p-value=0.0401; Input Resistance: p-value=0.6832; * p<0.05). (**C**) Average firing rate per current step in each condition. (Error bars represent SEM in all data panels).

**Figure S12.**
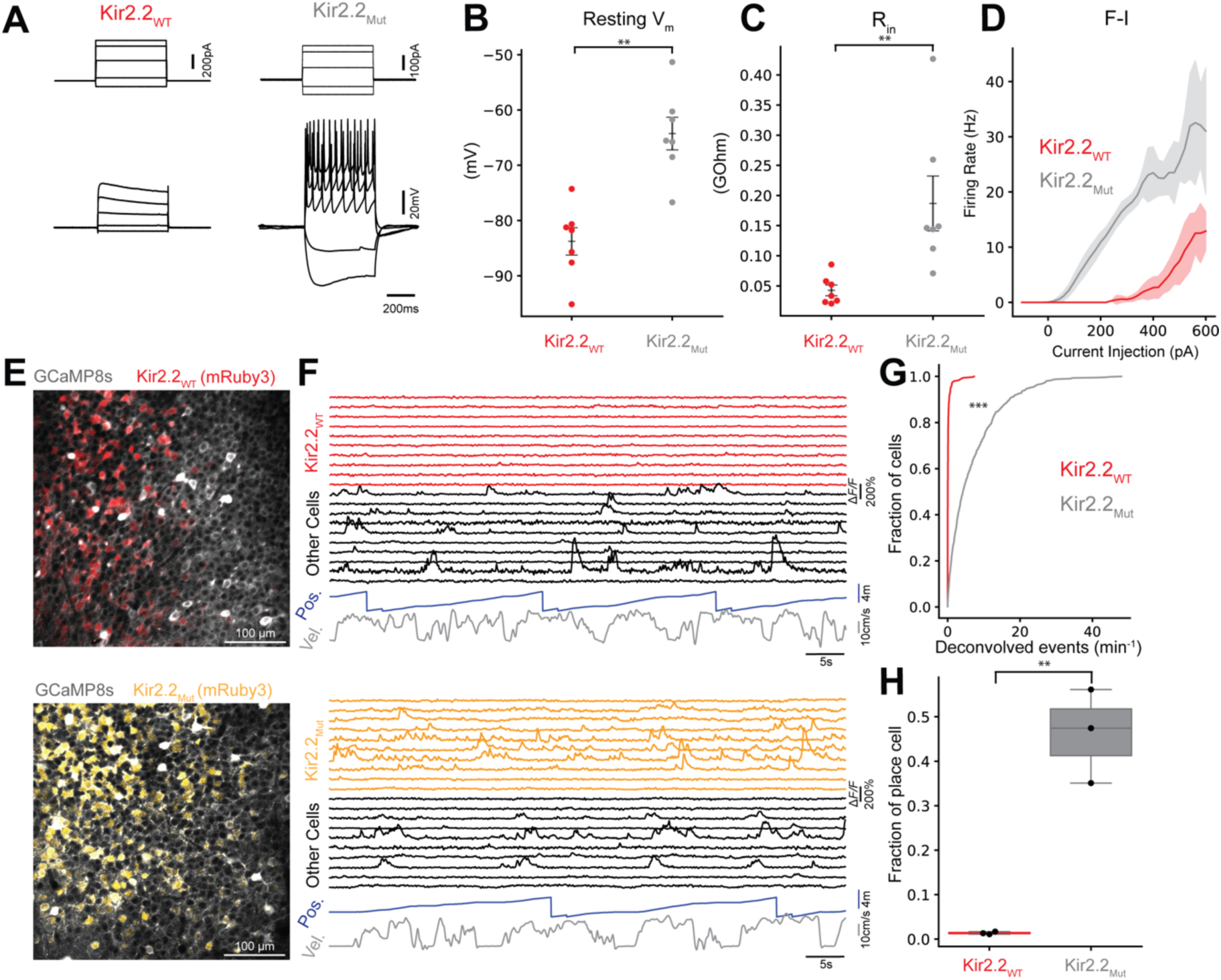
Ectopic expression of Kir2.2 silences CA1 neurons and reduces place cell percentage. (**A**) Representative traces of current (top) and voltage (bottom) as recorded in current clamp mode from cells that express either Kir2.2_WT_ or Kir2.2_Mut_. Cells that express Kir2.2_WT_ shows less action potentials despite higher current injections (current scale bar left: 200pA, right: 100pA). (**B,C**) Quantified data of resting membrane potential (**B**) and input resistance (**C**) in Kir2.2_WT_ and Kir2.2_Mut_ cells (Kir2.2_WT_ n = 7 cells from 2 mice; Kir2.2_Mut_ n = 7 cells from 2 mice). (Mann-Whitney U test. Resting Membrane Potential: p-value=0.001; Input Resistance: p-value=0.001). (**D**) Average firing rate per current step in each condition. (**E**) Representative 2P imaging field of view (FOV) of GCaMP and Kir2.2 channels in the CA1 pyramidal layer. Top: mouse is injected with AAV-hSyn-GCaMP8s, AAV-hSyn-FLEX-Kir2.2_WT_-mRuby3 and AAV-hSyn-Cre to express GCaMP and sparse expression of Kir2.2_WT_. Gray: GCaMP; Red: Kir2.2_WT_. Bottom: mouse is injected with AAV-hSyn-GCaMP8s, AAV-hSyn-FLEX-Kir2.2_Mut_-mRuby3 and AAV-hSyn-Cre to express GCaMP and sparse expression of Kir2.2_Mut_ as a control against Kir2.2_WT_. Gray: GCaMP; Orange: Kir2.2_Mut_.Scale bars: 100 mm. (**F**) Representative traces of dF/F, position and velocity of 2P recording during spatial navigation. For both top and bottom plots: top 10 traces are from cells that are infected with either Kir2.2_WT_ or Kir2.2_Mut_, bottom 10 traces are from cells in the same recording field of view but are not infected with either Kir2.2 AAVs. Note for overexpression of Kir2.2_WT_, other cells in the recording FOV are not affected by the channel overexpression and still exhibit normal calcium transients, and Kir2.2_Mut_ control cells still show normal activity. (**G**) Deconvolved events per minute from all cells across all mice that are infected with either Kir2.2 AAVs (n = 345 Kir2.2_WT_ cells from 3 mice, n = 620 Kir2.2_Mut_ cells from 3 mice, Mann-Whitney U Test, p-value = 1.74e-119). (**H**) Fraction of place cells in all cells from all mice that are infected with either Kir2.2 AAVs (Kir2.2_WT_ mean = 0.013, IQR = 0.003. Kir2.2_Mut_ mean = 0.462, IQR = 0.105. Unpaired t-test, p-value = 0.018). (Error bars represent SEM in all data panels).

**Figure S13.**
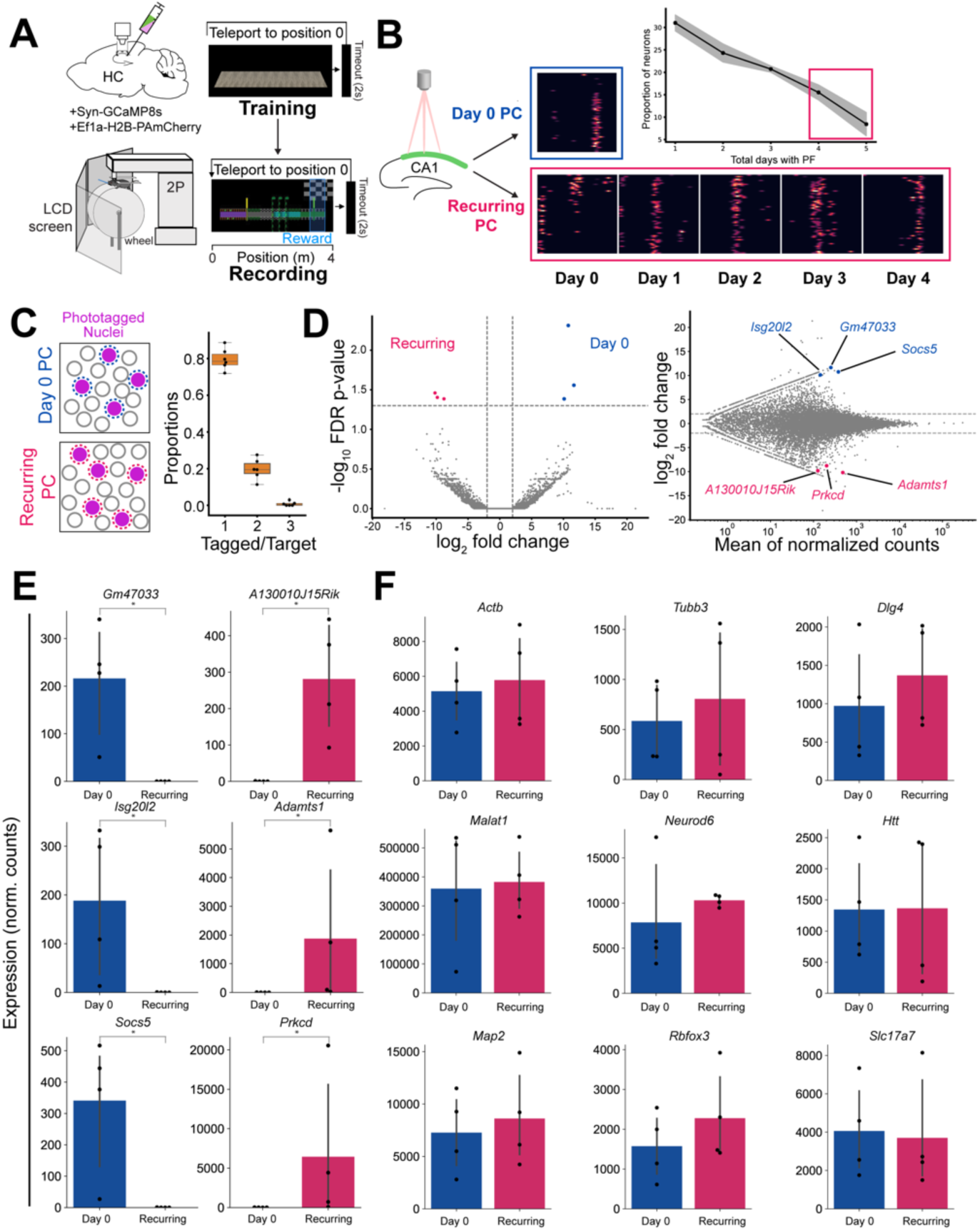
Transcriptional profiling of Day 0 and Recurring place cells. (**A**) Schematics of imaging and behavior setup. Mice were trained in featureless environments and recorded in a feature-rich environment. (**B**) Heatmaps of normalized ΔF/F activity from one sample day 0 place cell and one sample recurring place cell over 60 laps during VR navigation for either one day or across five days of recording. Upper right: quantification of proportion of neurons with respective numbers of place fields across 5 recording days (solid line: mean, shaded area: mean + s.e.m, n = 4 mice). (**C**) Left: schematics of *in vivo* photoactivated nuclei. ‘Day 0’ place cell sample and ‘Recurring’ place cell samples from different mice were collected for FACS and Meso-seq. Right: Proportion of single, double, and triple-tagged nuclei following phototagging of a single place cell. (**D**) Left: volcano plot of Meso-seq differential expressed gene (DEG) analysis for ‘Day0’ and ‘Recurring’ place cells. Right: Meso-seq MA plot depicting DeSeq2 normalized gene counts versus log2 fold change of Day 0/Recurring place cells samples. Day 0 and Recurring place cells are similar in genetic profiles with only 6 differentially expressed genes identified. Genes that are significantly different are labeled in blue and magenta (same as above). Genes shown in panels *E* are highlighted and labeled in *D*. (significantly different genes are shown in blue and magenta. blue: enriched in ‘Day 0’ cells; magenta: enriched in ‘Recurring’ cells). (**E**) Bar graph showing the normalized counts for differentially expressed genes (FDR adjusted p-value. *<0.05, \*\***<**0.01, ***<0.001, PyDeSeq2. Otherwise, comparisons are not significant). (**F**) Bar graph showing the normalized counts for genes that are not differentially expressed. Boxplots show the 25th, 50th (median), and 75th quartile ranges, with the whiskers extending to 1.5 interquartile ranges below or above the 25th or 75th quartiles, respectively. Outliers are defined as values extending beyond the whisker ranges.

**Supplementary Movie 1. Phototagging**

Real-time movie of *in vivo* two-photon imaging and phototagging of neurons. Imaging was performed at 1040 nm to visualize change in PAmCherry fluorescence. During phototagging, the 810-nm laser was scanned over target nuclei and the PMT was blanked. A frame average of 64 frames was applied for resolution and clarity. Final video was edited to include scale bar and laser switches. Video time is not representative of actual recording time.

**Supplementary Movie 2. Registered *in vivo* and *ex vivo* image stacks**

Three-dimensional rendering of registered z-stacks of *in vivo* and *ex vivo* tissue volume of the CA1 pyramidal layer with phototagged nuclei. Corresponding to *Figure 1F* and *Figure S1C,D*. Magenta: PAmCherry *in vivo* (captured on Bruker 2P microscope, wavelength = 1070 nm). Orange: PAmCherry *ex vivo* (captured on A1 HD25, Nikon Instruments Inc., wavelength = 568 nm)

**Supplementary Table 1. List of DEGs between place and silent cells**

List of DEGs between place and silent cells showing log 2-fold change, mean of normalized count (baseMean), and FDR adjusted p-value. All genes in the table have an FDR-adjusted p-value less than 0.05 when comparing silent versus place cells. log2FoldChange is computed for silent versus place (positive: enriched in silent cells, negative: enriched in place cells). File: Supplementary_Table.

